# Mouse and human microglial phenotypes in Alzheimer’s disease are controlled by amyloid plaque phagocytosis through Hif1α

**DOI:** 10.1101/639054

**Authors:** Alexandra Grubman, Xin Yi Choo, Gabriel Chew, John F. Ouyang, Guizhi Sun, Nathan P. Croft, Fernando J. Rossello, Rebecca Simmons, Sam Buckberry, Dulce Vargas Landin, Jahnvi Pflueger, Teresa H. Vandekolk, Zehra Abay, Xiaodong Liu, John M. Haynes, Catriona McLean, Sarah Williams, Siew Yeen Chai, Trevor Wilson, Ryan Lister, Colin W. Pouton, Anthony W. Purcell, Owen J. L. Rackham, Enrico Petretto, Jose M. Polo

## Abstract

The important role of microglia, the brain’s resident immune cells, in Alzheimer’s disease (AD) is now well recognized, however their molecular and functional diversity and underlying mechanisms still remain controversial. To transcriptionally and functionally characterize the diversity of microglia in AD and aging, we isolated the amyloid plaque-containing (XO4^+^) and non-containing (XO4^−^) microglia from an AD mouse model. Transcriptomics analysis unveiled independent transcriptional trajectories in ageing and AD. XO4^+^ microglial transcriptomes linked plaque phagocytosis to altered 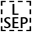 expression of *bona fide* late onset AD genetic risk factors. We further revealed that the XO4^+^ transcriptional program is present in a subset of human microglia from AD patients and is a direct and reversible consequence of Aβ plaque phagocytosis. Conversely, XO4^−^ microglia in AD displayed an accelerated ageing signature and contained more intracellular post synaptic material than plaque-containing microglia, despite reduced active synaptosome phagocytosis. Mechanistically, we predicted HIF1α as a core regulator of the XO4^−^/XO4^+^ axis, and further validated the mechanism *in vitro* using human stem cell-derived microglia like cells and primary human microglia. Together these findings unveiled the molecular mechanism underpinning the functional diversity of microglia in AD, providing opportunities to develop treatments targeted at subset specific manipulation of the microglial niche.

---

Microglia are specialist immune sentinel cells in the brain that respond to stranger or danger signals, remove cellular and extracellular debris, and regulate synaptic plasticity, maturation and removal^1–3^. Thus, microglial function is vital to physiological processes in the brain. AD is a progressive neurodegenerative condition, with no effective treatment options. Synapse loss, which occurs in cortical and hippocampal regions, most strongly correlates with cognitive dysfunction in AD^4^, and is accompanied by extracellular amyloid beta (Aβ) plaques and intraneuronal neurofibrillary tau tangles^5^. The possible role of microglia in AD has been highlighted by several unbiased data-driven functional genomics studies^6–11^. Indeed, all the identified genetic risk factors for the more common late onset AD (LOAD; >95% cases) have been recently reported to be microglia specific, or highly expressed in microglia^12^. Additionally, recent reports in mouse models, have found that microglia obtained from areas rich in plaques, have a different signature to microglia from plaque-free areas, however the origin of these microglia and their signature still remains unknown. Furthermore, knockouts of essential microglial receptor genes on an AD genetic background have yielded both protective and exacerbated phenotypes, often at different disease stages (i.e., ^13–17^ and ^18,19^). Coupled with the increasingly recognized spatio-temporal diversity of microglia^20–22^, these reports highlight the dynamic nature and complexity of microglial responses, which might be explained by the presence of multiple microglial subpopulations that may differentially affect disease course. To address this question, we used an aggressive plaque-depositing AD mouse to investigate whether differences in amyloid plaque phagocytosis are directly molecularly and functionally associated with specific microglia phenotypes in AD.

## RESULTS

### Methoxy-XO4 purifies molecularly distinct plaque-associated microglia populations

To understand the spatio-temporal and functional differences between plaque-phagocytic and non-phagocytic microglia in AD, we took advantage of *in vivo* Aβ plaque labeling using a fluorescent Congo-red derivative, methoxy-XO4^23^ (Fig. 1a-b, Extended data Fig. 1a-b). We found that 13.5% and 15.8% of cerebral microglia were amyloid-phagocytosing (XO4^+^) in 4, and 6m old 5xFAD mice, respectively, with no significant differences observed between male and female mice (Fig. 1c-d). Conversely, we did not detect any XO4^+^ microglia in 1m old 5xFAD mice, or WT mice of any age, and only 4.35% of cerebellar microglia in 5xFAD mice were XO4^+^, in accordance with the relative resistance of this region to pathology in mouse AD models and AD patients^24^.

**Figure 1:**
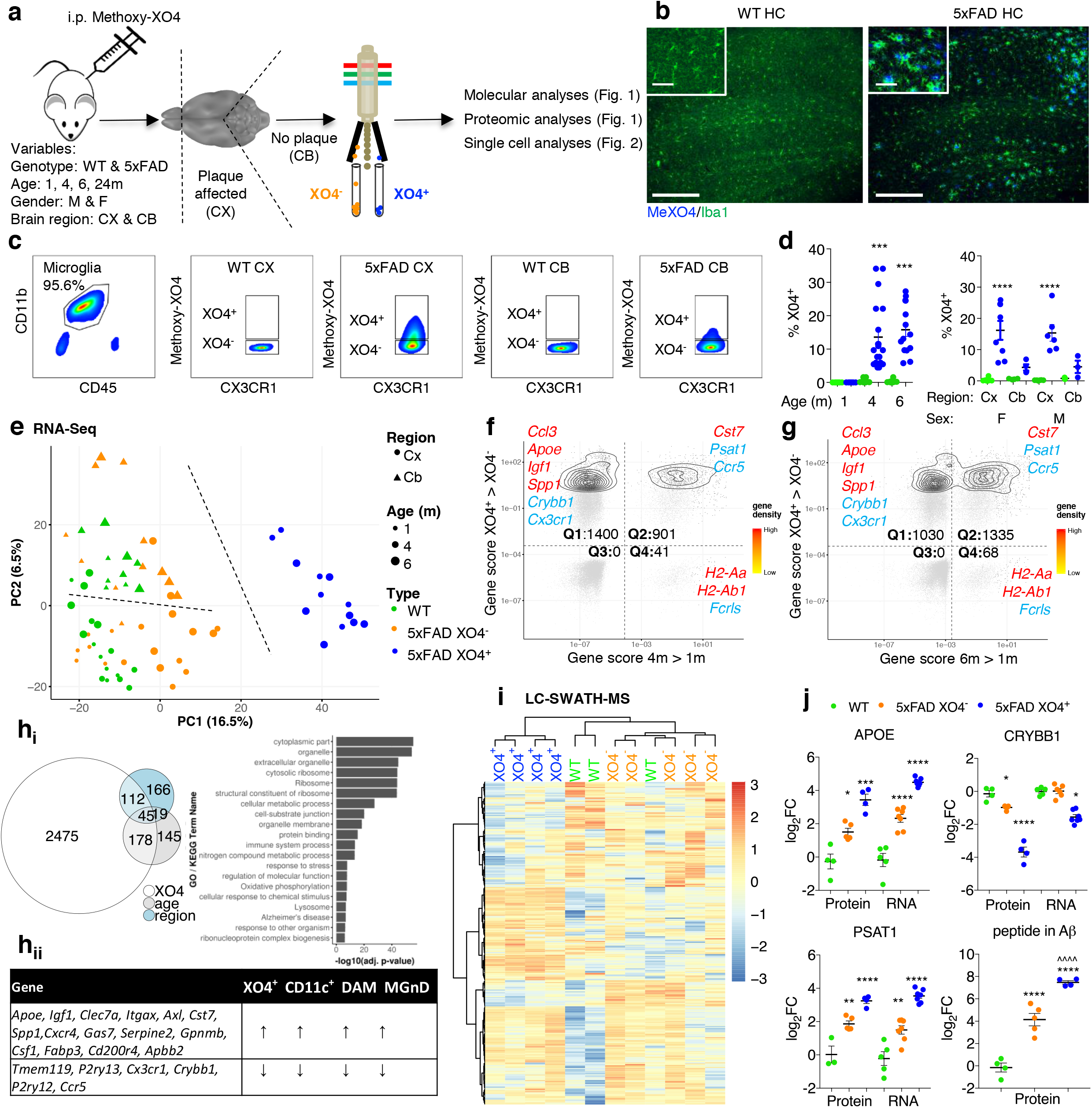
Methoxy-XO4 purifies a molecularly distinct plaque-phagocytic population in 5xFAD mice. **a**, Schematic of the methodology employed in this study**. b**, Representative immunofluorescence image of the hippocampus of WT and 5xFAD mice injected with Methoxy-XO4 and stained with Iba1 (AlexaFluor 488, *n*=6 animals per genotype), scale bar=250 μm, inset 50 μm **c**, Representative FACS plot (from *n*=12-19 animals per genotype and age group) showing that XO4^+^ microglia are present in 6m 5xFAD plaque-affected regions (top panels). **d**, left, the percentage of XO4^+^ microglia isolated from plaque-affected regions in 1, 4, and 6m old WT and 5xFAD mice, (*n*= 12-19 per genotype and age group; male and female mice pooled) and right, the percentage of XO4^+^ microglia isolated from plaque-affected and non-affected regions in 6m old male and female WT and 5xFAD mice (*n*= 6-8 per genotype), expressed as mean ± SEM. **e**, PCA of bulk RNA-seq. Cx, Cortex; Cb, Cerebellum **f, g** Gene cytometry plots showing genes that are differentially expressed between XO4^+^ and XO4^−^ microglia and/or genes that are differentially expressed between old (4, 6 month) and young (1 month) microglia. Gene scores are calculated as the product of the log fold change and –log_10_(FDR). Example genes in each quadrant are labelled in red (upregulated over time or phagocytosis) or blue (downregulated). **h_i_**, Venn diagram showing the overlap between genes whose expression levels could be explained by the age, region and XO4 covariate as well as GO and KEGG terms associated with XO4 covariate genes. **h_ii_**, table showing the 21 core microglial neurodegeneration signature genes and their direction of differential expression in DAM^28^, CD11c^+^ ^29^, MGnD^30^ and XO4^+^ microglia. **i**, heat map of targeted LC-SWATH-MS analysis of detected peptides within DEGs in *n*=3-5 biological replicates of WT (blue), XO4^−^ 5xFAD (orange) and XO4^+^ 5xFAD (green) microglia. **j**, comparison of RNA and protein expression for selected genes, and quantitation of a tryptic peptide in Aβ in microglia. Data are expressed as mean ± SEM log fold change compared to WT microglia, normalized relative to peptides in Supplementary table 2. *p* values in **d** and **j** were calculated by one-way ANOVA using Tukey’s multiple comparison test.

Next, we adapted CelSeq2 to transcriptionally profile a small number of XO4^+^ and XO4^−^ microglia during disease progression in 5xFAD mice^25^. XO4^−^ and XO4^+^ microglia clustered separately, showing a progression from WT to XO4^+^ microglia via XO4^−^ microglia in the first principal component (PC), whereas the second PC separated cerebellar microglia from their cerebral counterparts (Fig. 1e, Extended data Fig. 2a). We found that the factor explaining the greatest variance in gene expression was plaque phagocytosis (i.e., XO4^+^/XO4^−^, Extended data Fig. 2b). To investigate whether there was an interaction of plaque phagocytosis with age, we created “gene cytometry plots”, which show the distribution of differentially expressed (DE) genes by their significance score (i.e., false discovery rate (FDR)-weighted log fold change) for both age and XO4^+^/XO4^−^ factors from the DE analysis. These “gene cytometry plots” allowed us to identify gene sets that are significantly associated with either age, plaque internalization or both (Fig. 1f, g). Most DE genes (black points and contour plot) were associated with the XO4^+^ signature (Fig. 1f; Q1 and Q2) and detection of this signature was amplified by ageing, as 39% of XO4-associated genes were also associated with age progression in microglia at 4m (Fig. 1f; Q2), whereas this proportion had risen to 56% by 6m (Fig. 1g; Q2).

Overall, we identified 2,475 genes associated only with the plaque-phagocytic XO4^+^ state, which were enriched for the gene ontology (GO) terms ribosome, oxidative phosphorylation and phagolysosome pathways (Fig. 1h_i_, Supplementary table 1). These functional annotations suggest that the XO4^+^ gene signature reflects the microglia cell response to amyloid phagocytosis, requiring increased protein synthesis to enable efficient digestion of aggregated material, which is in keeping with the increased size of XO4^+^ compared to XO4^−^ microglia (Extended data Fig. 2c).

Among the most upregulated genes in XO4^+^ microglia were the two most highly penetrant late-onset AD risk factor genes, the receptor *Trem2* and its ligand *Apoe*^26^, as well as their interacting partners, *Tyrobp*^10^ and *Lpl, Ldlr, Lrpap1* (reviewed in ^27^), respectively, suggesting a link between phagocytosis and cholesterol transport pathways to the XO4^+^ phenotype in AD. Additionally, a significantly high proportion (63%, *p*=6.1×10^−11^ by hypergeometric test) of previously reported microglial sensome genes^28^ were differentially expressed in XO4^+^ microglia (Extended data Fig. 3a), including c-lectins (*Clec4a2, Clec4a3, Clec7a*) and CD markers (i.e., *Cd33, Cd68, Cd74, Cd180*). The XO4^+^ gene signature partially overlaps with recently reported microglia signatures obtained from APP/PS1 or 5xFAD model mice^29–31^ (Extended data Fig. 3b, c). Twenty-one core genes associated with XO4^+^/XO4^−^ phenotype were identified by all four studies^29–31^ as altered in neurodegenerative disease-associated microglia: *Apoe, Igf1, Clec7a, Itgax, Axl, Cst7, Spp1, Cxcr4, Gas7, Serpine2, Gpnmb, Csf1, Fabp3, Apbb2* and *Cd200r4* (upregulated in XO4^+^ microglia) and *Tmem119, P2ry13, Cx3cr1, Crybb1, P2ry12* and *Ccr5* (downregulated in XO4^+^ microglia; Fig. 1h_ii_). Notably, our analysis identified an additional 2,196 XO4^+^ signature genes that were not reported in any previous study (Extended data Fig. 3b). These newly identified genes were highly enriched for genes involved in oxidative phosphorylation, regulation of cell cycle, particularly M-phase (Extended data Fig. 3d) and immune response functions. Importantly, phagocytic associated genes were also highly enriched suggesting a potential role in augmented plaque clearance. Furthermore, XO4^+^ cells showed upregulation of *Mki67*, consistent with the observed expansion of the XO4^+^ subset during disease progression in 5xFAD mice. Alternatively, it is possible that non-plaque associated microglia proliferate and migrate towards plaques, as reported in APP/PS1 mice^32^.

This partial overlap and extended XO4^+^ network might be due to several factors: we profiled the specific microglia (CD11b^+^CD45^lo^CX3CR1^+^) population in AD that have phagocytosed plaques (XO4^+^), rather than more heterogeneous approaches (CD11c^+^ or Clec7a^+^) which did not include a marker for plaque *per se*, suggesting that the XO4^+^ microglia are an important, yet unappreciated, source of the different reported AD-associated signatures. Secondly, we were able to obtain a more homogenous dataset than either bulk APP/PS1 or CD11c^+^ microglia (which contain both plaque associated and distal microglia^29^), and we sequenced to a greater depth than the first study that defined the DAM microglia^29^.

To determine whether the observed transcriptional signature corresponded to changes in the proteome of XO4^+^ microglia, we performed targeted LC-SWATH-MS analysis (Fig. 1i). Out of 174 proteins corresponding to DEGs detected, downregulation of 19 genes in XO4^+^ microglia was mirrored at the protein level, including for DOCK10, CRYBB1, PLXNA1 and FGD2. We also detected elevated concentrations of peptides corresponding to 99 genes upregulated in XO4^+^ microglia, including APOE, PSAT1, RPS4X, ANXA5, GAPDH, LGALSBP, CD180, C1QB and LRPAP1. Nonetheless, we also observed several instances where transcripts induced in XO4^+^ microglia were repressed post-transcriptionally (SERPINE2, ACACA, UQCRH, and MYO5A). We next sought to directly validate the presence of Aβ within XO4^+^ cells. In 5xFAD, but not WT, microglia, we detected a single peptide within the amyloid precursor protein sequence (LVFFAEDVGSNK), compared to multiple tryptic peptides arising from APP fragments that were detected within purified synaptosome fractions (Supplementary table 2). That LVFFAEDVGSNK is the only tryptic peptide present within the Aβ sequence suggests that this peptide arose from microglial phagocytosis Aβ and not from microglial expression of *App* or phagocytosis of APP. The 180-fold upregulation of the Aβ peptide in XO4^+^ microglia compared to the limit of detection further confirmed that these microglia contain high levels of internalized Aβ (Fig. 1j). Surprisingly, XO4^−^ microglia also contained Aβ, albeit at ~10 fold lower levels than XO4^+^ microglia. This is consistent with low levels of non-fibrillar (i.e., oligomeric) Aβ phagocytosis by XO4^−^ microglia, although our observation could also be explained by low numbers of contaminating XO4^+^ cells containing partially digested Aβ peptide fragments and thus not detected by Methoxy-XO4 staining. Together these data show that the molecular state of XO4^+^ microglia is distinct from homeostatic and XO4^−^ microglia at the RNA and protein levels and is directly associated with high levels of internalized fibrillar Aβ.

### Two distinct molecular processes drive microglial alterations in AD

Recent reports highlight subtle transcriptional differences between individual microglial cells, even those residing within the same anatomical regions^33,34^. Thus, to examine the heterogeneity inside the XO4^+^ cell population, we first employed viSNE^35^ on FACS data sets, a dimensionality reduction and visualization approach that preserves multi-variate relationships within single-cell data. We labelled myeloid cells with a subset of 6 microglial sensome markers^28^ including the genetic LOAD risk factors CD33^36^ and TREM2^8^ that regulate microglial Aβ phagocytosis^16,37^. XO4^+^ labeled microglia clustered together, and revealed uniformly high expression of CD33, TREM2 and the Csf1 receptor, CD115 (Fig. 2a, Extended data Fig. 4a; *p<*0.0001 compared to XO4^−^ microglia for each marker by 2-tailed paired *t*-test), consistent with the reported increase of CD33-expressing microglia in human AD^37^ and TREM2 upregulation in microglial processes directly interacting with plaques in three AD mouse models^16^. It is noteworthy that while all microglia homogeneously expressed CD11b, CX3CR1 and low levels of CD45, the expression pattern of the archetypal microglia-specific proteins, CD115, CD33 and TREM2 were highly variable in individual WT and 5xFAD XO4^−^ microglia. Thus, we further investigated microglial heterogeneity by single cell sequencing using the 10X Genomics Chromium system. As aging is the most important risk factor for LOAD^38^, and microglia are known to express an altered ageing gene signature^20,39^, we examined whether microglial subpopulations in 5xFAD mice adopted an aging phenotype by including FACS-sorted microglia from WT adult (6m) and WT old mice (24m) as well as both XO4^−^ and XO4^+^ populations from a 6m old 5xFAD mouse (Fig. 2b). Similar to our bulk analyses of XO4^+/−^ microglia, the first PC was dominated by the shift to a phagocytic phenotype, whereas the third PC separated 6m from aged WT cells, while 5xFAD XO4^−^ cells were also shifted in the direction of aged WT microglia (Fig. 2b). Single-Cell Consensus Clustering (SC3)^40^ separated microglia into four clusters, whereby all 5xFAD XO4^+^ cells were entirely contained within cluster 2, and cluster 3 was almost entirely composed of WT 6m microglia (Fig. 2c). Interestingly, 6m XO4^−^ microglia and aged WT microglia clustered together (i.e., have similar transcriptional profiles), suggesting that the XO4^−^ population, from AD mice, may represent an accelerated aging phenotype. The ageing signature was enriched in α-defensin genes (Fig. 2d_i-ii_), antimicrobial peptides which mobilize immune cells and enhance phagocytosis in the periphery (reviewed in ^41^), although their role in the brain has not previously been described. In agreement with our bulk RNA-seq data (Fig. 1e-h), we observed loss of homeostatic genes (i.e., *Crybb1*, Fig. 2d_iii_) and upregulation of XO4^+^ signature genes (i.e., *Igf1, Ccl3*, Fig. 2d_iv-v_) in the XO4^+^ microglial population.

**Figure 2:**
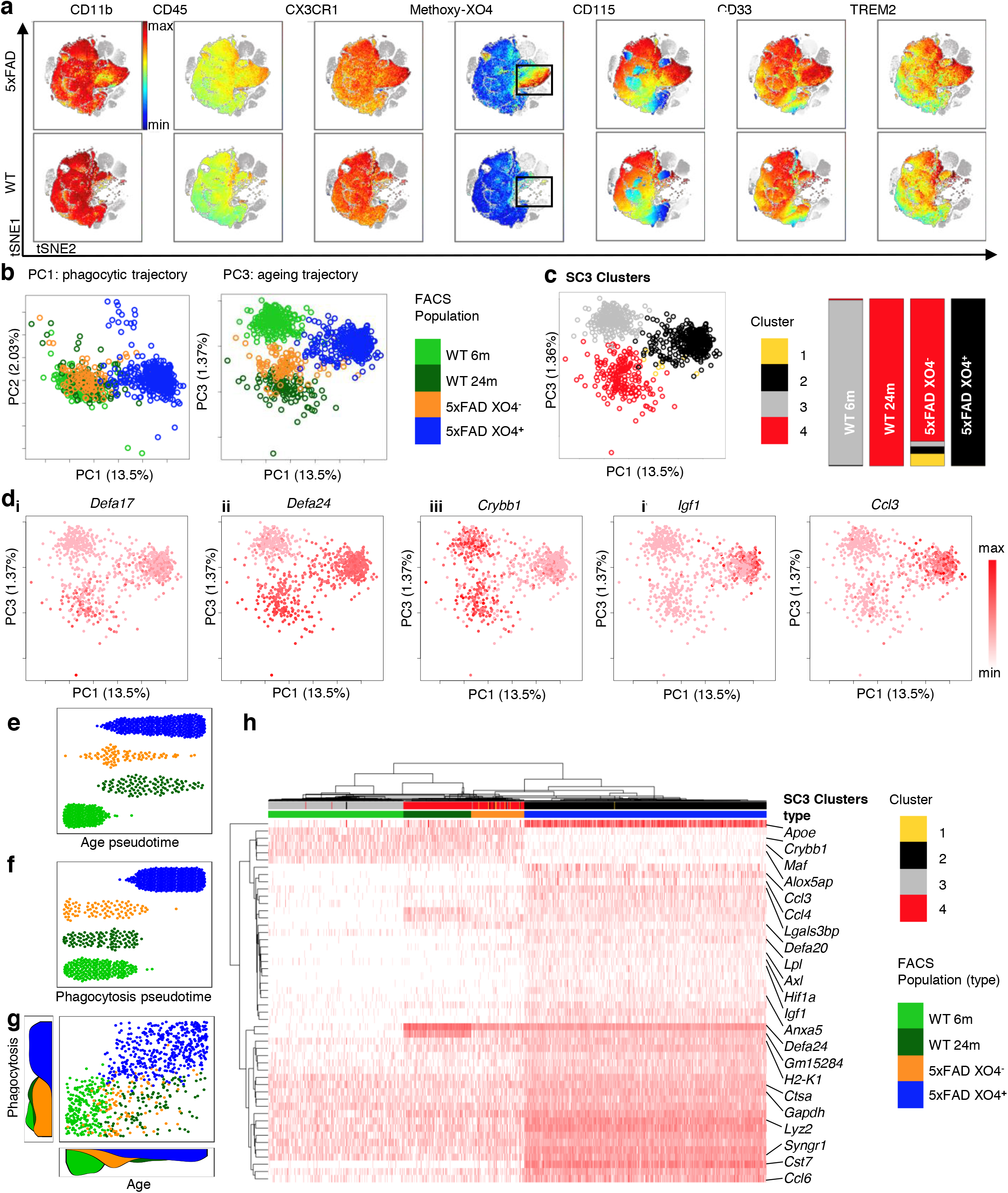
Single cell sequencing identifies an ageing profile in 5xFAD XO4^−^ microglia. **a**, Dimensionality reduction representation (viSNE, representative of *n*=3 mice per genotype) of myeloid cells isolated from WT (top) and 5xFAD (bottom) 6m male mice. Microglia (CD11b^+^CD45^lo^CX3CR1^+^) are colored for expression of CD11b, CD45, CX3CR1, Methoxy-XO4, CD115, CD33 and TREM2, whereas remaining myeloid cells are grayscale for clarity. **b**, PCA of 893 single cells x 1671 feature genes showing the distribution of cells from each FACS-sorted sample. **c**, PCA plot of single microglia colored by SC3 clusters and composition of automated clusters as a percentage of sequenced FACS-sorted cell populations. **d**. PCA plots for single microglia colored by expression of selected ageing microglia genes (i-ii), homeostatic (iii) and XO4^+^ signature genes (iv-v). **e, f** Diffusion maps pseudotime analysis of microglial populations ordered by their expression of **e**, ageing DEGs (6m WT v 24m WT, 42 DEGs) or **f**, phagocytic DEGs (6m 5xFAD XO4^−^ v 6m 5xFAD XO4^+^, 474 DEGs) **g**, scatter plot showing the relationship between ageing and phagocytosing pseudotime in individual cells, and the density of cells at each point during the ageing (bottom) and phagocytosing (left) trajectories. **h**, Hierarchical clustering and heat map showing expression of the top 50 DEGs across the 4 SC3 clusters.

To further elucidate the molecular microglial trajectories in 5xFAD mice, we ordered cells in aging pseudotime, which we defined by their expression of the 42 ageing-specific DEGs (FDR<0.05, top 20 genes reported in Extended data Fig. 5a, list in Supplementary table 3) between adult and aged WT microglia (Fig. 2e). This analysis indicated that the XO4^−^ and XO4^+^ groups were both shifted in the ageing trajectory, although there was a high degree of heterogeneity of the pseudoage of individual cells within 5xFAD microglial populations, suggesting a gradual acquisition of the ageing signature. Conversely, when cells were ordered by their phagocytic pseudotime, taking into account the 474 phagocytosis-specific DEGs (FDR<0.05, top 20 genes reported in Extended data Fig. 5b, list in Supplementary table 3) between phagocytosing and non-phagocytosing AD microglia, we observed a profile that resembled a switch from a non-phagocytic to a phagocytic signature with few cells exhibiting an intermediate or transitional phagocytic pseudotime (Fig. 2f). These data suggest that although the XO4^+^ and XO4^−^ molecular signatures are distinct, the cellular age of individual microglia lies on a continuum within each 5xFAD population and is controlled by an independent component of ageing, which is largely unrelated to phagocytosis (Fig. 2g). The top 50 most variable genes in the dataset included microglial identity genes (*i.e. Crybb1, Alox5ap, Maf*), α-defensin genes (*Defa20, Defa21, Defa24, Gm15284*), chemokines (*Ccl3, Ccl4, Ccl6*) and lysosomal genes (*Lyz2, Cst7, Ctsa*), and clustered samples according to both age and phagocytic phenotype (Fig. 2h). In addition to the newly described set of genes (356 XO4^+^ specific genes not identified in DAM microglia), our single cell RNA-seq data in XO4^+^ cells also recapitulate core Stage I and Stage II changes described for DAM microglia^29^. However, the XO4^−^ and aged microglia molecular signatures do not overlap with either Stage I or II DAM (Extended data Fig. 6a, b), although they do differ from homeostatic microglia (Fig. 2b-g). These data collectively suggest that the XO4^−^ microglia are on an independent trajectory and are not necessary cellular intermediates poised to become XO4^+^ cells.

### The XO4^+^ molecular signature is reversible and can be induced by phagocytosis of amyloid plaques

We next asked whether the XO4^+^ transcriptional program is a consequence of AD brain microenvironment *per se* or if the phagocytosis of plaques is the signal required for this molecular switch. In other words, we wished to address whether the XO4^+^ signature predates plaque phagocytosis and enables enhanced phagocytic capacity of a proportion of cells in 5xFAD mice or is a result of plaque phagocytosis. Thus, to determine whether and how any microglia could activate an XO4^+^ signature, we cultured CFSE-labelled WT microglia with organotypic hippocampal slice cultures (OHSCs) from 6m 5xFAD mice (Fig. 3a). To ensure that the XO4^+^ microglial signature was not dependent on the presence of the Methoxy-XO4 dye, we instead labelled OHSCs with NIAD4^42^, an alternative fluorescent Aβ-binding dye. To establish that a healthy and plaque-associated microglial signature can be detected using this system, we also cultured WT microglia in WT OHSCs and 5xFAD microglia in 5xFAD OHSCs, respectively. We FACS sorted groups of 10 endogenous (CSFE^−^) and exogenous (CFSE^+^) microglial cells that were plaque positive (NIAD4^+^) or not (NIAD4^−^) for molecular profiling. qPCR analysis of a panel of 42 homeostatic and XO4^+^ signature genes identified two main clusters of microglia cells by SC3 (Fig. 3b, Supplementary table 4). Cluster 1 contained all but 1 sample of NIAD4^+^ microglia, including both endogenous 5xFAD cells that had phagocytosed plaques, and exogenous (CFSE^+^) cells from either WT or 5xFAD animals that had acquired the XO4^+^ signature upon phagocytosis of plaques in 5xFAD OHSCs (Fig. 3d, e). These microglia downregulated numerous homeostatic microglia signature genes (*Cx3cr1, Maf, P2ry12*, and *Cebpa;* Fig. 3c_i-ii_, Extended data Fig. 7a), and activated expression of XO4^+^ profile genes (*Cst7, Igf1*, *Apoe, Spp1*, *Lgals3, Trem2, Lyz2*, and *Cstd;* Fig. 3c_iii-iv_, Extended data Fig. 7b). Therefore, these data not only demonstrate that the XO4^+^ signature is gained following phagocytosis of plaques but also demonstrates that both WT and 5xFAD microglia are capable of acquiring the XO4^+^ signature Furthermore, most WT cells cultured with 5xFAD OHSCs that had not phagocytosed plaques retained their homeostatic signature (Fig. 3e, *p*=0.0326 comparing population proportions), suggesting that microglial plaque phagocytosis, and not mere exposure to an AD-like brain microenvironment is the trigger for conversion to the XO4^+^ state. Conversely, cluster 2 was characterized by high expression of homeostatic microglial genes and low expression of XO4^+^ microglia signature markers and included all microglial cells isolated from WT OHSCs. This suggests that a physiological environment is not capable of sustaining an XO4^+^ signature. Whether this occurs through reversal of the XO4^+^ signature upon lysosomal degradation of internalized plaque or via increased susceptibility to cell death of XO4^+^ cells in a homeostatic environment will be important to address in future studies. Our data are more consistent with the reversal of XO4^+^ signature in a physiological environment, as over half of exogenous CFSE^+^ 5xFAD cells isolated from 5xFAD OHSCs molecularly resembled XO4^+^ microglia (Fig. 3f) suggesting that survival is not impacted, whereas all 5xFAD CFSE^+^ cells recovered from WT OHSCs clustered with homeostatic microglia (Fig. 3g, *p*=0.0155). Interestingly, phagocytosis of synaptosome-labelled beads by microglia in OHSCs did not induce Apoe or Trem2 expression, consistent with the XO4+ signature resulting specifically from phagocytosis of plaques rather than the phagocytic process per se (Extended data Fig. 7c). Collectively, our data show that the XO4^+^ program is activated by plaque phagocytosis and is reversible by exposure to a healthy brain microenvironment.

**Figure 3:**
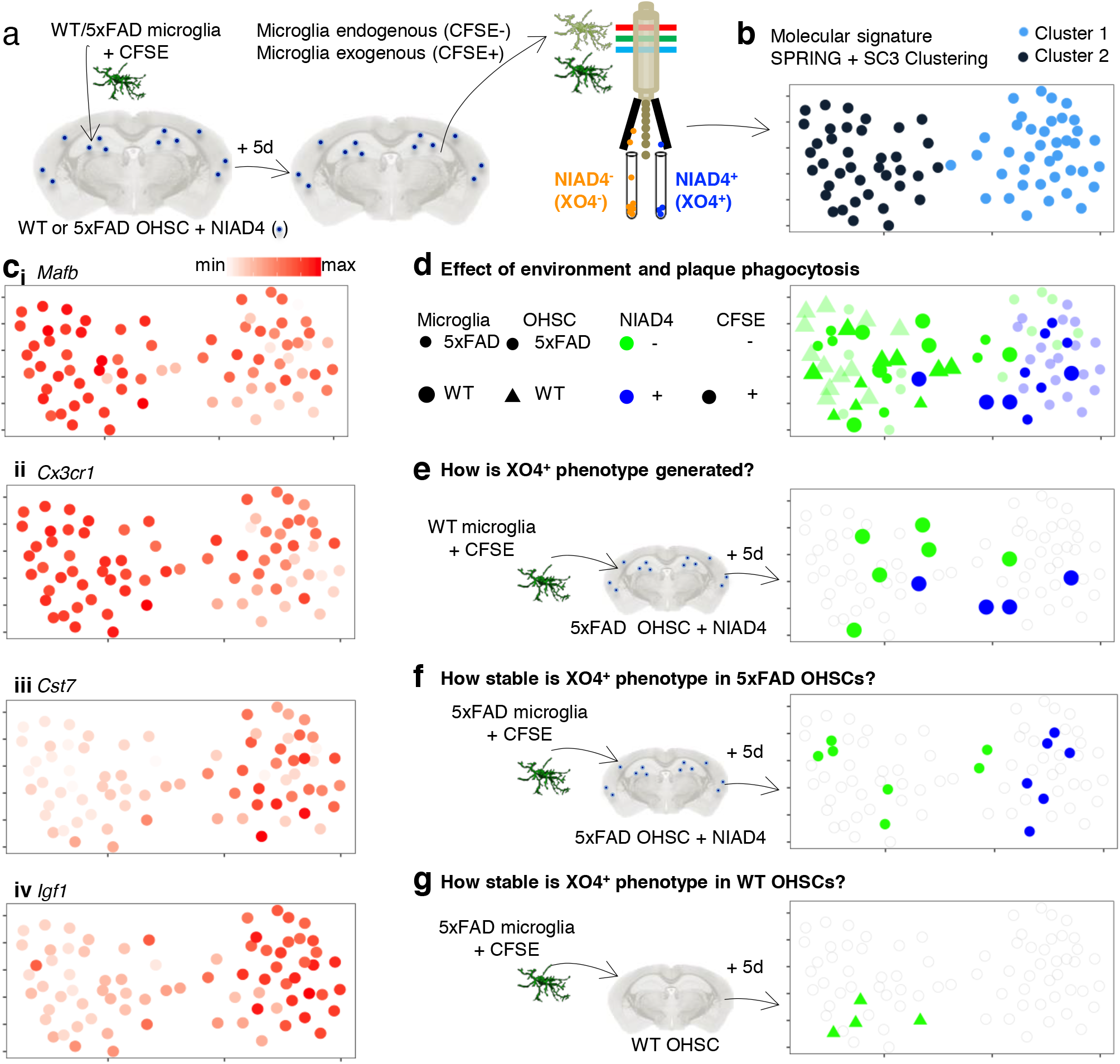
The XO4^+^ molecular signature is reversible and is acquired through phagocytosis of amyloid plaques. **a**, Schematic representing the experimental design to FACS isolate CFSE-labelled replenished and CFSE^−^ endogenous microglia that differentially phagocytose endogenous NIAD4-labelled plaques after 5 days co-culture with WT or 5xFAD OHSCs. **b-d**, k-nearest-neighbor graph rendered using a force directed layout^103^, colored by SC3 cluster (**b**), and log2 transformed ΔCt values of selected DEGs (**c**) or cell grouping (**d**). **e**, WT microglia added into 5xFAD slices recapitulate a XO4^+^ signature upon plaque phagocytosis (*p*=0.0326). **f**, XO4^+^ phenotype is stable in exogenous CFSE^+^NIAD4^+^ 5xFAD microglia recovered from 5xFAD slices, but **g**, is lost in CFSE^+^ 5xFAD microglia recovered from WT slices (*p*=0.0155).

### XO4^+^ microglia actively internalize more synaptosomes *ex vivo*

Aβ oligomers have been reported to induce microglia to aberrantly engulf synapses via deregulated complement deposition^43^, although it is currently unknown whether all microglia equally participate in this process. Thus, we hypothesized that akin to their transcriptional differences as a consequence of phagocytosis of plaques, XO4^+^ and XO4^−^ microglia in AD may also differentially engulf synaptic proteins. To study this, we first visualized and quantified internalization of the post-synaptic marker, PSD95^44^ by individual microglia in the dentate gyrus of WT and 5xFAD mice (Fig. 4a, Extended data Fig. 8a). There was a significant increase in steady-state internal PSD95 by XO4^−^ microglia compared to XO4^+^ microglia in 6m old 5xFAD mice (Fig. 4b). Given the previously reported susceptibility of excitatory cholinergic neurons and network dysfunction in inhibitory neurons in AD^45^, and to examine whether steady state levels of pre-synaptic components internalized within microglia differed between XO4^+^ and XO4^−^ microglia, we examined GAD65 and found that there were also less GAD65^+^ synapses within XO4^+^ microglia (Fig. 4c,d). Given recent data demonstrating that synapse pruning is a complement-dependent process^43,46^, we examined colocalization of C3 deposition with microglia in 5xFAD mice and observed decreased complement C3 co-localization with XO4^+^ microglia compared to XO4^−^ cells (Fig. 4e, f). Indeed, both at the transcriptional and protein level, we detected higher levels of complement components in XO4^−^ microglia than in XO4^+^ microglia (Extended data Fig. 8e).

**Figure 4:**
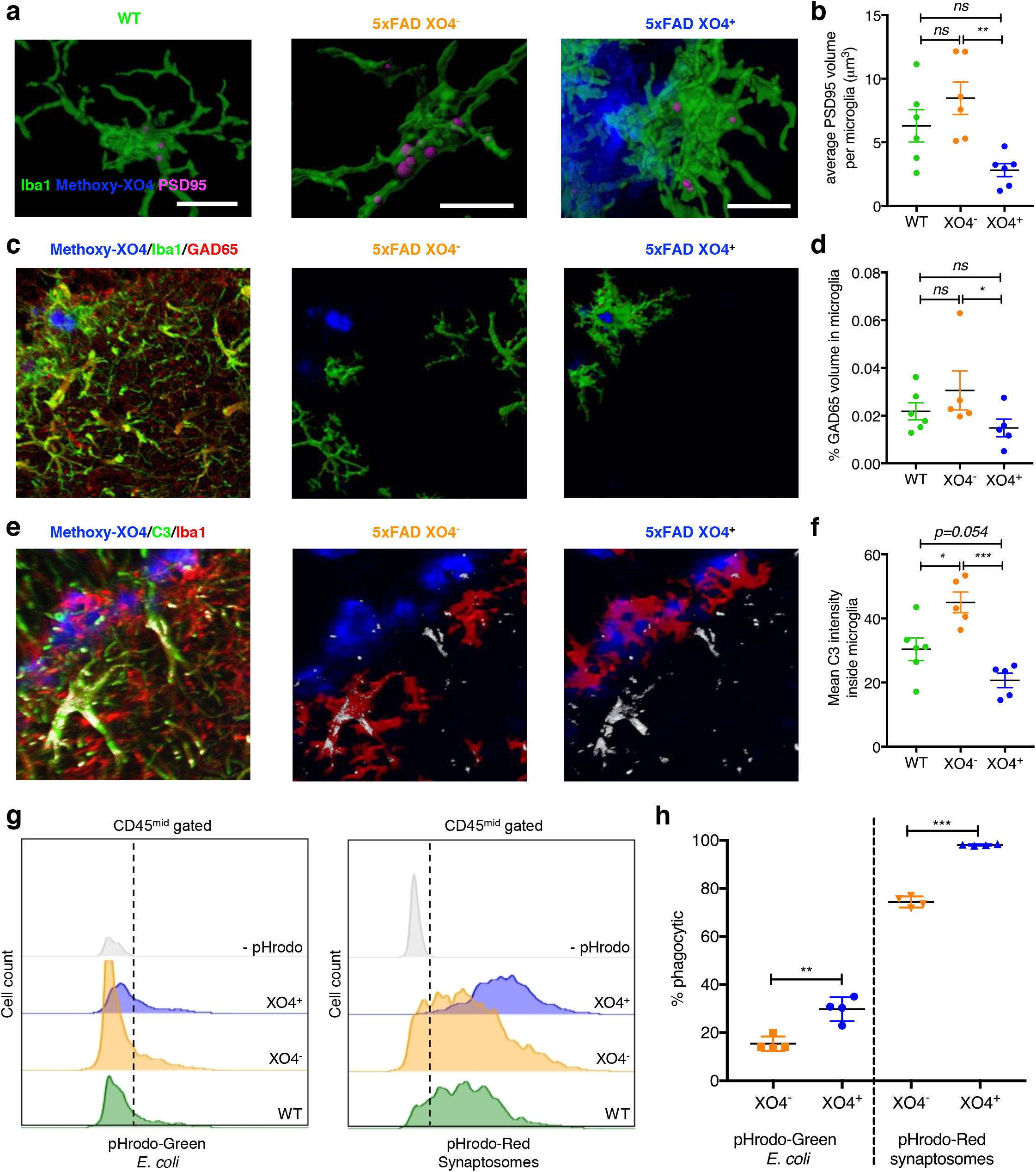
5xFAD XO4^+^ microglia contain less post synaptic material than 5xFAD XO4^−^ microglia in the dentate gyrus. **a, c, e**, Representative 3D reconstructions of confocal z-stacks showing PSD95 (**a**), GAD95 (**c**) or complement C3 (**e**) internalized within WT, 5xFAD XO4^−^ or 5xFAD XO4^+^ microglia cells (scale bars = 15 μm). **b**, PSD95 within microglia quantified as the average volume of phagocytosed PSD95 volume per microglia volume in each dentate gyrus or cerebellar section (*n* =6 z-stacks per condition; \**p* < 0.01, using 2 tailed *t-*test) **d**, GAD65 within microglia quantified as the % volume of phagocytosed GAD65 in microglia (*n* =5 z-stacks per condition; \**p* < 0.01, using 2 tailed *t-*test. Note that removal of the outlier sample does not increase the *p* value. **f**, Mean C3 intensity within microglia (*n* =5 z-stacks per condition; \**p* < 0.01, using 2 tailed *t-*test). All data is from *n*=3-6 WT and *n*=5-6 5xFAD animals and is presented as mean ± SEM. **g.** Functional analysis of *ex vivo* mouse microglia phagocytosis following 1 h incubation with pHrodo-red labelled synaptosomes or pHrodo-green labelled *E. coli* by FACS. Each population is gated based on XO4^+^ signal and compared to controls not incubated with pHrodo particles. h. Quantitation of the percentage of XO4^+^ and XO4^−^ microglia that phagocytose pHrodo-red labelled synaptosomes or pHrodo-green labelled *E. coli.* \*\**p* < 0.001 and \*\*\**p* < 0.0001 by paired 2-tailed *t*-test.

Confocal measurements are static by nature and thus do not distinguish between differences in flux through the phagolysosomal pathway, and observed differences could thus result from altered internalization rate, rate of degradation and efflux. Furthermore, the relative difference in synaptic densities near a plaque (lower in plaque core than >30 μm from the plaque halo (Extended data Fig. 8b), consistent with previous studies^47^), makes the interpretation of this static measurement difficult. Therefore, in order to directly examine active internalization of synaptic material by microglia, we performed functional phagocytosis assays with pHrodo-red labelled synaptosomes on freshly isolated *ex vivo* microglia from 5xFAD mice (Fig. 4g, h). We detected a higher rate of internalization by XO4^+^ microglia by FACS. pHrodo-green labelled *E. coli* particles were also more efficiently phagocytosed by XO4^+^ microglia compared to their XO4^−^ counterparts (Fig. 4g, h), suggesting that XO4^+^ microglia are primed for increased phagocytic capacity, in accordance with their increased expression of lysosomal machinery (Fig. 1h). In summary, these data suggest that, in addition to their distinct molecular profiles, as a result of amyloid plaque phagocytosis the microglial subsets in the hippocampus of 5xFAD mice exhibit different rates of synapse internalization.

### Human AD microglia recapitulate the XO4^+^ microglia signature

Imaging data show that plaque-adjacent microglia in human post mortem AD brains upregulate LPL^29^ and downregulate P2RY12^31^, consistent with an XO4^+^ signature surrounding plaques. However, no study has yet profiled microglial subsets in brain tissue from AD patients. Thus, we used fluorescence activated nuclear sorting (FANS)^48^ and DroNc-Seq^49^, to obtain the first single cell transcriptomes of human brain AD patients (*n*=6) and neurologically normal controls (*n*=6) (see companion resource paper; Grubman *et al*. 2019). Focusing on the microglia cell sub-population, Uniform Manifold Approximation and Projection (UMAP) analysis, a visualization tool for single cell data, clearly separated AD from healthy cells (Fig. 5a). We further used these data to test whether the XO4^+^ gene signature identified in 5xFAD mice is conserved in human AD. We took the top 10% of DEGs (*n*=167) between 5xFAD XO4^+^ and XO4^−^ microglia to represent the XO4^+^ signature. After getting the human orthologues of the mouse DEGs, we recapitulated the XO4^+^ pseudotime signature in AD patients (Fig. 5b, d, Supplementary table 5). In keeping with the mouse data (Fig. 2f), 63.4% of the human AD microglia sub-population exhibited a significant switch from a non-phagocytic to a phagocytic signature (defined as >50th percentile of the phagocytosis pseudotime; *p*=6.82×10^−6^, hypergeometric test), whereas only 41.9% of healthy cells exhibited the phagocytic signature (*p*=1, hypergeometric test). Similarly, human cells ordered by the mouse aging pseudotime gene signature showed a significant association with AD (Fig. 5c, e, *p*=0.00484, hypergeometric test) but not control microglia (*p*=0.997, hypergeometric test). Interestingly, the association between AD microglia and the ageing signature was primarily driven by genes overlapping with the XO4^+^ signature, as the association was no longer significant when only genes specific to the ageing signature were analyzed (Extended data Fig. 9a, *n*=128 genes, *p*=0.476). The GWAS hit gene *FRMD4A* and the homeostatic microglia gene, *CX3CR1*, were downregulated specifically in the subset of human AD microglia with high XO4^+^ scores, whereas *LINGO1* and *CRYAB* were specifically upregulated in this subset (Fig. 5g). Furthermore, there is a significant preservation of mouse XO4^+^ signature in human AD patients (Fig. 5h; *p*=6.41×10^−4^). Interestingly, the TREM2-adaptor *TYROBP*, and *APOE* were upregulated in microglia with high XO4^+^ scores, irrespective of the origin of microglia from control or AD patients, suggesting that there are AD-specific and common components of this signature (Fig. 5g). Altogether, our results confirm the existence of multiple subsets of microglia in human AD, including a subset that maps to the XO4^+^ mouse signature identified here, and an increased ageing signature primarily driven by the overlap with the XO4^+^ signature.

**Figure 5.**
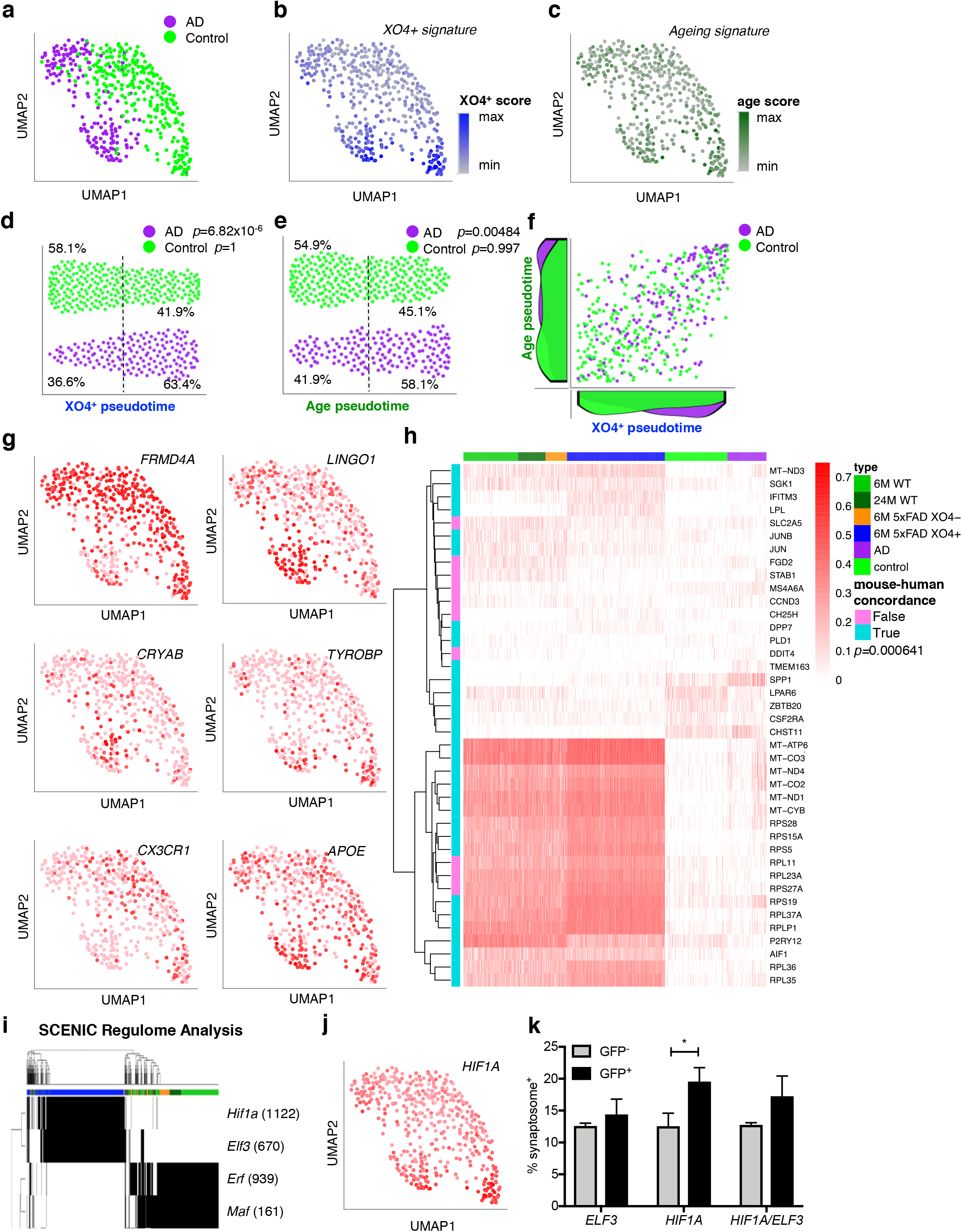
The XO4^+^ signature is molecularly and functionally replicated in human AD microglia. **a-c** UMAP projection of single microglia nuclei from control and AD patient frontal cortex, cases (*n*=172 nuclei) and controls (*n*=277 nuclei). The microglial population was determined by similarity to known microglial markers^95^. The UMAP projection is colored by **a**, disease diagnosis **b**, XO4^+^ score **c**, or ageing signature score. Diffusion maps pseudotime analysis of microglial populations ordered by their expression of **d**, XO4^+^ DEGs (taking top 10% of respective DE genes regardless of overlap with aging DEGs, 167 DEGs between 5xFAD XO4^+^ and XO4^−^ mice) or **e**, ageing DEGs (top 10% or 167 DEGs between 24M WT and 6M WT mice). **f**, scatter plot showing the relationship between ageing and XO4^+^ pseudotime in individual cells and the density of cells at each point during the ageing (left) and XO4^+^ (bottom) trajectories. **g**, UMAP projection of single human microglia colored by expression of selected cluster specific-genes. **h** Hierarchical clustering and heat map showing expression of the overlapping DEGs in each of the 4 mouse microglia clusters with the DEGs between human control and AD microglia. The mouse-human concordance on the direction of change between control and AD (human) or WT and XO4^+^ populations is shown for each gene (*p*=0.000641). **i**, SCENIC regulon analysis showing that *Hif1a* and *Elf3* are predicted to control the XO4^+^ gene regulatory network. **j**, UMAP projection of single human microglia colored by *HIF1A* regulon activity. **k**, Fluorescently labeled synaptosome internalization by primary microglia transfected with GFP-tagged inducible *HIF1A* and/or *ELF3* overexpression constructs. The data are presented as mean ± SEM and show the difference in synaptosome internalization between GFP^+^ and GFP^−^ (non-transfected) cells tested from within the same well.

We next stained human sections with 6E10 to label plaques, PSD95 and Iba1 and calculated semi-quantitatively the amount of post-synaptic material in plaque-adjacent and distal microglia (Extended data Fig. 9b-d). Analogous to our findings in 5xFAD mice, we observed that plaque-associated microglia in human AD patients contained modestly, but significantly (*p*=0.02), less PSD95 staining than in microglia present in the same brain region but not associated with plaques (Extended data Fig. 9d). To functionally assess the effect of the XO4^+^ identity on synapse phagocytosis in human cells, and due to the impossibility of isolating *bona fide* methoxy-XO4 stained *human* microglia, we first needed to understand the gene regulatory network(s) underlying the XO4^+^ signature, to mimic it *in vitro.* To this end, we used the Single Cell Regulatory Network Inference and Clustering (SCENIC)^50^ method; briefly, this approach identifies *regulons* (i.e., group of genes that are controlled by a common regulatory gene) by grouping the predicted target genes of a given transcription factor (TF) module that show motif enrichment for the same TF (see Methods for details). In addition, SCENIC infers the “activity” of the *regulon* in each cell by the AUCell algorithm, providing a way to associate the regulon activity to a specific cell population (e.g., XO4^+^). SCENIC analysis identified the *Hif1a* and *Elf3* regulons with the highest regulon activity in XO4^+^ cells (Fig. 5i, Supplementary table 6), and we found *HIF1A* regulon activity to be upregulated in the human microglial nuclei that displayed a high XO4^+^ score (Fig. 5j). These two regulons had minimal overlap of downstream regulated genes, however, functional enrichment analysis identified that both *Hif1a* and *Elf3* specific networks captured many of the same functional processes, including ribosome, oxidative phosphorylation, Alzheimer’s disease and metabolic pathways (Extended data Fig. 10a-b). We thus examined whether *HIF1A* and/or *ELF3* overexpression could contribute to priming human microglia towards an overactive synaptosome-phagocytic phenotype. We transfected primary human microglia with a dox-inducible GFP-tagged construct to genetically turn on these regulomes, and found that *HIF1A* overexpression indeed functionally increased synaptosome phagocytosis, consistent with a role for *HIF1A* in regulating XO4^+^ functions (Fig. 5k). Together, our data show that the XO4^+^ signature is recapitulated in a subset of human AD microglia and can be controlled via *HIF1A* to increase synaptosome phagocytosis.

### *HIF1A* regulon controls the XO4^+^ molecular signature through MYD88 and mTOR

To examine potential upstream small molecule targets and downstream readouts of the XO4^+^ network and *HIF1A* regulome, we employed Ingenuity Pathway Analysis (IPA)^51^. IPA identified several upstream regulators of the *Hif1a* regulon (Fig. 6a, Supplementary table 7), including BMP9 (also known as GDF2), a ligand of the transforming growth factor-β (TGF-β) superfamily, the Toll like receptor (TLR) adaptor MyD88 and mTOR, which were previously involved in restricting amyloidosis^52^, microglial response to pathogens^53^ and pro-inflammatory microglia^54^ (Fig. 6b). As our data suggest that the XO4^+^ molecular and functional properties extend to human AD (Fig. 5), we chose to validate these regulons in human ESC-derived microglia-like cells (iMGLs), using a recently published protocol^55^. Stimulation of iMGLs with the MyD88-signalling dependent TLR1/2 agonist, Pam3csk, alone, or in combination with BMP9, but not with the MyD88-independent TLR3 agonist, Poly:IC, resulted in upregulation of *HIF1A* mRNA (Fig. 6c). We also noted induction of downstream IPA-predicted targets *SPP1* and *GAPDH*, as well as secretion of chemokines MIP1α and MIPβ, encoded by the XO4^+^ signature genes, *CCL3* and *CCL4*, respectively (Fig. 6c, d; Extended data Fig. 10c). However, an XO4^+^ signature gene that was not part of the *Hif1a* regulon, *TREM2*, was repressed by Pam3csk (Extended data Fig. 10c). Despite this, we found by FACS that TREM2 protein on the iMGL surface was slightly upregulated by Pam3Csk and BMP9, indicating that stimulation of part of the network may drive downstream changes to induce the XO4^+^ signature (Extended data Fig. 10d). As predicted by IPA, treatment of cells with the mTOR inhibitor, rapamycin, blocked MyD88-and BMP9-dependent induction of *Hif1a* regulon XO4^+^ signature mRNA expression (Fig. 6e; Extended data Fig. 10e). Moreover, rapamycin treatment reduced TREM2 expression at both the RNA and protein levels and induced the homeostatic marker CX3CR1 (Extended data Fig. 10c-d). Indeed, the network of genes induced by Pam3csk and repressed by rapamycin significantly overlapped with the XO4^+^ signature (64 genes, *p*=3.17×10^−20^ by hypergeometric test), and was enriched for gene ontologies including lysosome, *HIF1A*, and lipid metabolism (Fig. 6f, Extended data Fig. 10e). Moreover, rapamycin was able to reduce synaptosome phagocytosis in iMGLs treated with *in vitro*-formed Aβ fibrils (Fig. 6g). Together these data show not only the predicted regulatory role of the *Hif1α* regulon on the XO4^+^ signature (Fig. 5i-j), but that components of this regulon can be modulated by upstream small molecules to control microglia cell fate along the homeostatic to XO4^+^ signature axis.

**Figure 6.**
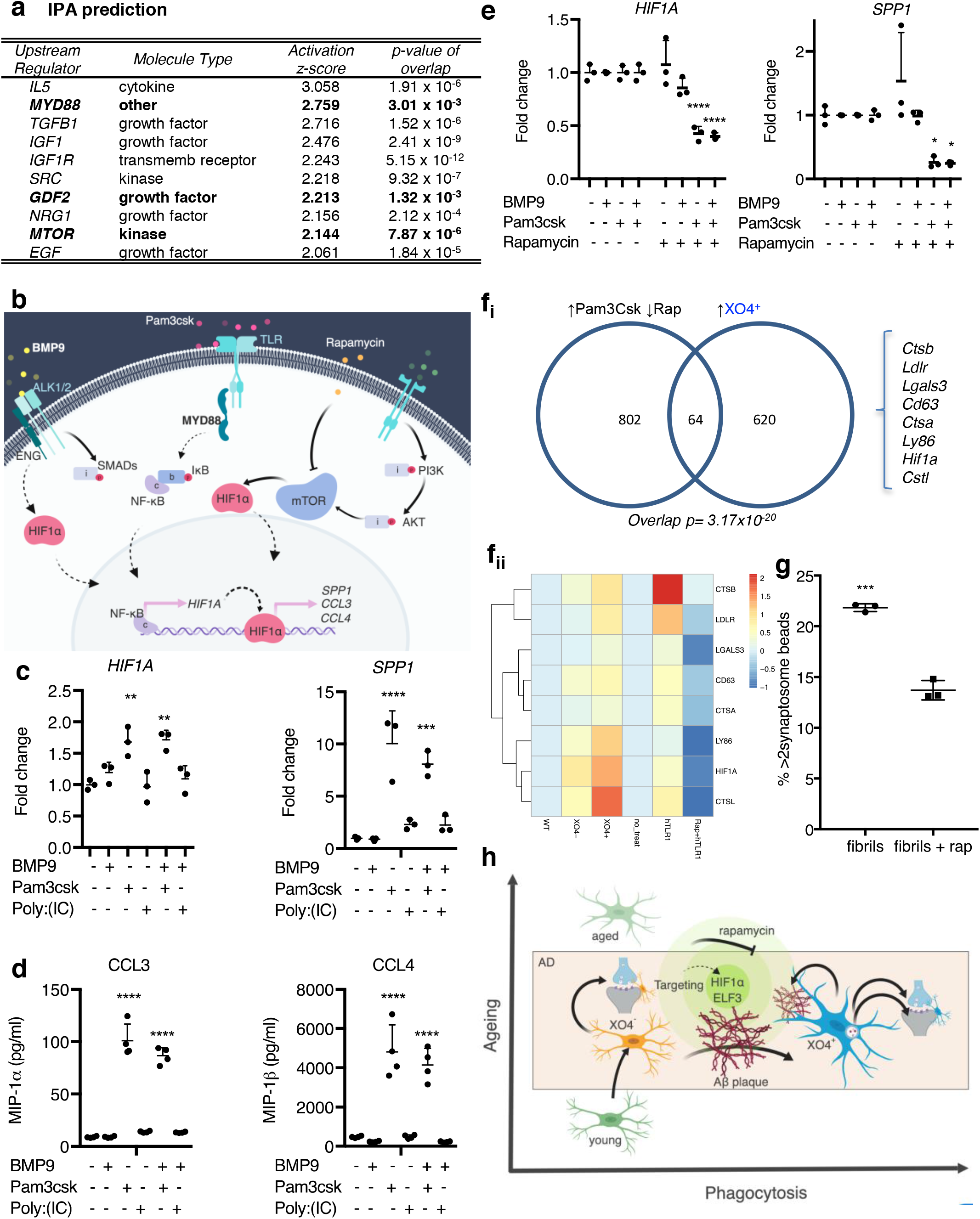
Microglial XO4^+^ signature can be manipulated through the *Hif1a* regulon. **a**, Top ten activators of the *Hif1a* regulon predicted by IPA. The activation z-score is a statistical measure of the match between the expected relationship direction of regulation and the observed gene expression; positive z-scores are indicative of predicted activation. *p*-value of overlap refers to the significance of the overlap between the *Hif1a* regulon gene set and the regulated target genes predicted by IPA. **b**, Cartoon diagram of hypothesis generated by IPA. **c**, Stimulation of iMGLs with MyD88-dependent TLR agonist Pam3csk (alone or with BMP9) induces XO4^+^ signature genes within the *Hif1a* regulon as identified by qPCR and **d**, cytometric bead array. MyD88-independent TLR stimulation (Poly:IC) does not shift iMGLs towards an XO4^+^ signature. Data are fold changes normalized to non-treated cells. **e**, MyD88-dependent XO4^+^ signature gene expression is modulated by rapamycin. Data are fold changes induced by rapamycin normalized to each respective treatment in the absence of rapamycin. *p* values throughout are calculated by one way ANOVA and Sidak post-test. **f**, Venn diagram showing the overlap between a XO4^+^-like state induced in iMGLs using Pam3csk and reversed by rapamycin (RNA-seq) as predicted by ingenuity pathway analysis (IPA) with the mouse XO4^+^ signature, as measured by RNA-seq. Representative gene expression heatmap of the overlap, showing expression levels in mice (WT, 5xFAD XO4^−^, 5xFAD XO4^+^) and human iMGLs (non-treated, Pam3csk and Pam3csk+rapamycin). **g**, Fluorescently labeled synaptosome internalization by iMGLs treated with amyloid fibrils, alone, or in combination with rapamycin for 48 h, as measured by FACS. The data are presented as mean ± SEM, and analysed by unpaired *t-*test. **h**, proposed model of generation and regulation of microglia diversity in AD.

## DISCUSSION

Two recent seminal publications^29,31^ identified a novel microglial phenotype near Aβ plaques that was dependent on *Trem2* and *Apoe* in two mouse models of AD, the transgenic APP/PS1 and 5xFAD strains that overexpress mutant human *APP* and *PSEN1* specifically in neurons. In our study, we chose the aggressive 5xFAD model of AD for several reasons: firstly, it allowed us to decouple the ageing component of AD from amyloid phagocytosis because pathology in these mice occurs prior to ageing. Secondly, synapse loss in this model facilitated investigation of subset-specific microglial synapse pruning functions. Here, we substantially expanded on previous work by firstly defining the specific signature in a subset of AD microglia as a result of plaque phagocytosis *in vivo* and defined the mechanisms responsible for induction and maintenance of this phenotype. More critically, we found the same signature in a subset of human AD microglia and started to address the controversy regarding the beneficial^56^ or detrimental^31^ functional role of these cells. A key challenge to tackling this question had resulted from the lack of a robust method to isolate these cells directly. Keren-Shaul *et. al.* addressed this problem through single cell transcriptomics, which, although allowing them to distinguish the cells, did not permit additional functional characterizations due to the destructive nature of the technique. Kraseman *et. al.* chose instead to purify these cells through either sorting for CLEC7A, a marker that was also present on a subset of WT microglia, or by purifying microglia that phagocytosed apoptotic neurons injected stereotaxically into mice. Here, we chose to take up this challenge using a direct approach with methoxy-XO4 staining^23^, combined with both single cell, bulk transcriptome and proteomic analyses followed by functional characterization of synapse internalization.

Our findings show that two distinct but interrelated processes drive microglial changes in AD: accelerated aging, as well as a direct response to plaque phagocytosis. Keren-Shaul reported that 3% of aged WT microglia exhibited a DAM signature that was undetected in younger animals. As the DAM population is not defined by the functional uptake of amyloid plaque and hence may collectively be comprised of both X04^+^ and X04^−^ microglia, the effects of ageing and amyloid phagocytosis on microglial transcriptional changes could not be assessed independently in that study. In comparison, our data shows that the age-associated signature acquired by XO4^−^ microglia is independent of uptake of amyloid plaque, thus allowing us to disentangle the ageing from amyloid phagocytic processes. The transcriptional signature of aged human microglia has been reported by two groups previously. Post mortem microglia from cognitively normal patients displayed similar gene expression profiles to mouse microglia, but human and mouse signatures diverged significantly with ageing^57^. The second human dataset (HuMi_Aged) was enriched for susceptibility genes for LOAD^39^, which is consistent with our mouse data, despite the aggressive nature of our EOAD model. Several recent studies suggested changes to microglia gene expression in human AD, however were not able to interrogate the specific cell subsets responsible for this signature as they were performed in tissue rather than single cell or single nuclei as per our study. For example, AD-specific gene upregulation of *TREM2, TYROBP, CLEC7A, CD68, CD34*, *SPP1* and various MHC Class II genes were described^10,58^ and ThioS (plaque associated)-positive microglia in 4/5 human AD patients tested also stained positive for LPL^29^. In addition, Yin *et al*. recently reported increased *APOE, AXL, TREM2* and *HLA-DRA* mRNA expression in laser microdissected dense-core plaque-adjacent regions in the medial frontal gyrus compared to plaque distal regions from the same sections in EOAD patients, although the same phenomenon was not observed in LOAD patients^59^. Thus, our study sheds light into an elusive cellular population unveiling the *bona fide* microglial heterogeneity and their transcriptional make up in AD patients. It is worthwhile noting, that the mouse XO4^+^ signature is present in 5xFAD mice under conditions of plaque deposition with little tau pathology^60^, whereas human AD pathology invariably includes tau, or may present with additional pathological phenotypes which likely produce altered signatures and responses in microglia. Interestingly, we observe that the microglial cells from human AD patients that cluster with control microglia are mostly composed of cells from female patient brains. While the current study lacks the power to conclude that female microglia may be impaired at mounting XO4^+^ responses, these findings warrant a more in depth (and larger) study focused on contribution of gender to transcriptional variability of microglia, particularly in the context of gender differences in AD incidence^61^.

The results presented in this study extend the repertoire of microglia types beyond the DAM classification. Our data are consistent with XO4^−^ microglia cells being a case of functional deregulation, set on a trajectory of accelerated ageing and not a transcriptional intermediate *en route* to become X04^+^, and therefore different from what was reported for Stage I and II DAM. XO4^−^ contain more steady-state internal synaptic material than XO4^+^ microglia (Fig. 4a-d) despite their reduced capacity for active phagocytosis (Fig. 4g-h), do not upregulate TREM2 (Fig. 1f-g) and do not migrate towards plaques, despite some capacity to internalize amyloid (Fig. 1j). This hypothesis fits well with current theories regarding the synaptotoxic role of oligomers, and suggests that XO4^−^ microglia internalize oligomeric Aβ, which is found enriched throughout 5xFAD brain except regions containing and directly surrounding fibrillar Aβ plaques^62^. Whether the toxic role of XO4^−^ microglia and the ageing signature they acquire is a direct result of oligomer phagocytosis remains to be investigated. On the other hand, plaque phagocytosis resulting in a XO4^+^ signature primes microglia for efficient phagocytosis of multiple substrates including synaptosomes. Rapid pruning of damaged synapses near dystrophic neurites localized around plaques, appears to be, at least initially, protective and may go awry later in disease progression as described before^63^. Our results are consistent with recent studies, showing improved behavior in AD mouse models that was associated with enhanced microglial amyloid plaque phagocytosis in response to scanning ultrasound^64^, and IL-33 treatment^65^. Indeed, IL-33 intracellular signaling occurs exclusively via MyD88, which is consistent with our prediction that MyD88-dependent mechanisms regulate the transition from XO4^−^ to XO4^+^.

We showed that all microglia possess an innate capacity to activate the XO4^+^ signature, and our data show that phagocytosis of amyloid plaque *per se* is necessary and sufficient for the generation of this disease associated transcriptomic signature. Interestingly unlike the partially overlapping MGnD signature reported by Krasemann *et al* 2017, the XO4^+^ microglial signature does not appear to be caused by phagocytosis of neurons. If the XO4^+^ signature was induced by phagocytosis of dying neurons, a proportion of microglia isolated from WT OHCSs would convert to an XO4^+^ signature, as OHSC culture from adult animals inevitably results in some neuron death. However, the XO4^+^ signature is only observed in WT microglia that have been cultured with an AD OHSC and phagocytosed NIAD4-labelled fibrillar amyloid, and not in microglia that have phagocytosed synaptosome-conjugated pHrodo particles.

The rapid switch in the transcriptomic profile of microglia is supported by our pseudotime analyses and demonstrates the remarkable plasticity of microglia. Although the receptors involved in recognition of synapses are distinct from those responsible for amyloid recognition^3,43,66^, consistent with our data, there is evidence that Aβ and synapse engulfment may be inhibited together by C3 knockout in APP/PS1 mice^67^, or enhanced together by conditional microglia specific knockout of TDP-43^44^. We uncovered that the *Hif1a* regulon in part underlies the molecular mechanism for a transition to the XO4^+^ phenotype and regulates synaptosome phagocytosis, and importantly, a portion of this network can be manipulated *in vitro* as predicted by our IPA, through combined stimulation with BMP9 and MyD88-dependent signaling pathways, and can be partially reversed by mTOR blockade via rapamycin. On this note, our demonstration of this pathway in iMGLs opens new avenues to examine how and to what extent the patient’s genetic background may influence the manipulation of the XO4^+/−^ axis via *HIF1A* in patient-derived iMGLs. A *Hif1a* epigenetic and transcriptomic signature was recently identified in microglia following immune training by peripheral LPS administration in APP/PS1 mice^68^. There was little gene overlap between the *Hif1a* module in ^68^ and the *Hif1a* regulon identified here, reinforcing the importance of microglial fine tuning of context-dependent responses to specific stimuli. Targeting MyD88 alone has yielded contradictory effects in AD models^69,70^, which could possibly result from differential effects on XO4^+^ and XO4^−^ populations. Thus, modulation of MyD88-dependent pro-inflammatory responses by combination with the protective effects of BMP9 on cognitive function^52,71^ may cover a larger part of the gene regulatory network to induce conversion of XO4^−^ cells into XO4^+^. As yet, while we provide a first description of the gene regulatory network, the upstream regulators controlling *TREM2* and *APOE* in this network remain to be determined.

We hypothesize a model whereby microglia during ageing are set on a transcriptional trajectory which is accelerated in AD microenvironment, however upon plaque phagocytosis those microglia re-route into a different trajectory through a Hif1α regulon, resulting in efficient phagocytosis of synaptic components around plaques. We show how the microglial gene networks we uncovered can be harnessed by computational prediction of microglial subset-targeting drugs, or via network pharmacology and repositioning approaches. In view of the apparent beneficial roles of XO4^+^ microglia, potential therapeutic strategies could involve the targeted conversion of XO4^−^ to XO4^+^ microglia by using small molecules to activate the right transcriptional networks.

## METHODS

### Animals

Heterozygous 5xFAD transgenic mice (B6SJL hybrid background) over-expressing FAD mutant forms of human APP (Swedish mutation K670/ 671NL, London mutation V717I, and Florida mutation I716V) and PSEN1 (M146L and L286V), regulated by the neuron-specific mouse *Thy1* promoter^60^ were housed at Monash Animal Research Platform (MARP) under specific pathogen-free conditions in a day-night controlled light cycle, provided with food and water *ad libitum*. Animals were used for experiments at different ages through adulthood, as indicated, without undergoing any procedures prior to their stated use. All use and handling of animals for experimentation was approved by Monash Animal Ethics Committee (MARP/2016/112) and conformed to national and institutional guidelines.

### Human patient demographics

Paraffin embedded human frontal cortex sections of post-mortem Alzheimer’s disease and non-disease aged matched individuals (10 μm) were obtained from the Victorian Brain Bank (Ethics Approval: MUHREC 2016-0554; patient demographics in table below).

| Case # | Age | Gender | PMI | Diagnosis |
| --- | --- | --- | --- | --- |
| 03/965 | 59 | Female | 30 | Control |
| 05/854 | 63.4 | Female | 30.5 | Control |
| V13/020 | 65.8 | Female | 43 | Control |
| 08/026 | 67.3 | Female | 24 | Control |
| 04/157 | 68.3 | Female | 71.5 | Control |
| 07/239 | 78.8 | Female | 19 | Control |
| V14/008 | 79 | Female | 32 | Control |
| 06/993 | 80.7 | Female | 59 | Control |
| V12/019 | 81.2 | Female | 25 | Control |
| 07/143 | 82.7 | Female | 28.5 | Control |
| V15/013 | 62.5 | Female | 5.5 | AD |
| V12/009 | 64.7 | Female | 21 | AD |
| V11/046 | 65.9 | Female | 40 | AD |
| V15/064 | 67.8 | Female | 21 | AD |
| V16/031 | 68.4 | Female | 31 | AD |
| 04/042 | 78.9 | Female | 19.5 | AD |
| V16/043 | 79.9 | Female | 35 | AD |
| V15/027 | 80.3 | Female | 62 | AD |
| V13/004 | 82.5 | Female | 15.5 | AD |
| V12/038 | 82.7 | Female | 8 | AD |

### Nuclei isolation sorting from human AD brain tissue

Nuclei were prepared for single nucleus RNA-seq from post mortem frozen Entorhinal Cortex of control individuals and AD patients (patient demographics in table below). Nuclei isolation was carried out using the Nuclei Isolation Kit: Nuclei EZ Prep (Sigma, #NUC101) as described^49^. Briefly, tissue samples were homogenized using a glass dounce grinder in 2 ml of ice-cold EZ PREP and incubated on ice for 5 min. Centrifuged nuclei (500 × g, 5 min and 4 °C) were washed in ice-cold EZ PREP buffer, and Nuclei Suspension Buffer (NSB; consisting of 1× PBS, 1% (w/v) BSA and 0.2 U/μl RNase inhibitor (Clontech, #2313A). Isolated nuclei were resuspended in NSB to 10^6^ nuclei per 400 μl), filtered through a 40 μm cell strainer and counted with Trypan blue. Nuclei enriched in Nuclei Suspension Buffer were stained with DAPI (1:1000) for nuclei isolation using the FACSAria™ III cell sorter (BD Biosciences, Franklin Lakes, NJ; 70 μm nozzle, 21-22 psi). Nuclei were defined as DAPI^+^ singlets. Sorted nuclei were counted twice loaded onto the 10X Chromium (10X Genomics). Library construction was performed using the Chromium Single Cell 3′ Library & Gel Bead Kit v2 (10X Genomics, #PN-120237) with 18 cDNA pre-amplification cycles and sequencing using high-output lanes of the NextSeq 500 (Illumina).

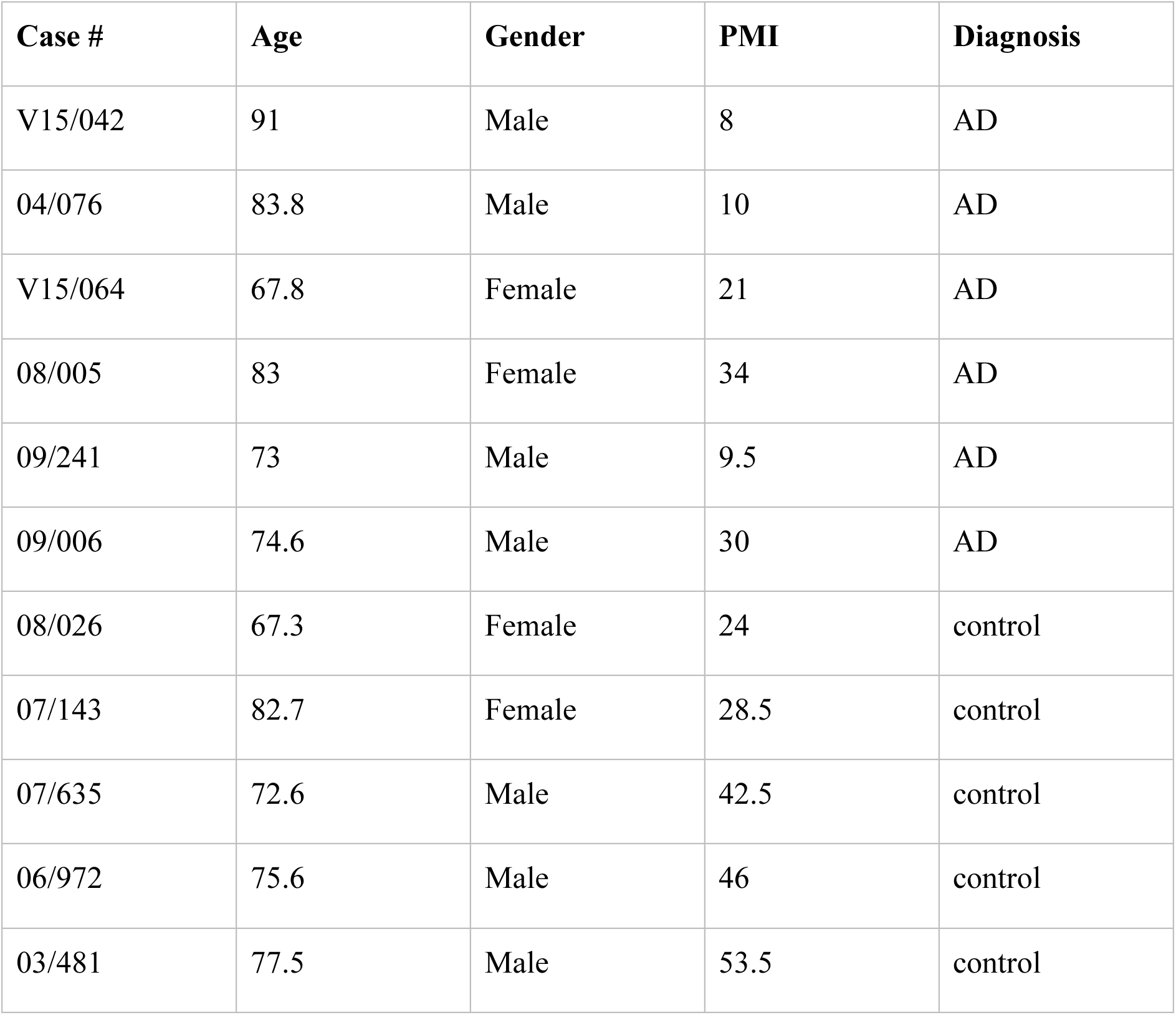

| Case # | Age | Gender | PMI | Diagnosis |
| --- | --- | --- | --- | --- |
| V15/042 | 91 | Male | 8 | AD |
| 04/076 | 83.8 | Male | 10 | AD |
| V15/064 | 67.8 | Female | 21 | AD |
| 08/005 | 83 | Female | 34 | AD |
| 09/241 | 73 | Male | 9.5 | AD |
| 09/006 | 74.6 | Male | 30 | AD |
| 08/026 | 67.3 | Female | 24 | control |
| 07/143 | 82.7 | Female | 28.5 | control |
| 07/635 | 72.6 | Male | 42.5 | control |
| 06/972 | 75.6 | Male | 46 | control |
| 03/481 | 77.5 | Male | 53.5 | control |

### Acute isolation of microglia and fluorescence activated cell sorting

2 h prior to killing, mice were injected intraperitoneally with Methoxy–X04 (2 mg/ml in 1:1 ratio of DMSO to 0.9% (w/v) NaCl, pH 12) at 5 mg/kg. Mice were euthanized by CO_2_ and transcardially perfused with ice-cold PBS prior to brain extraction. Whole brains, excluding brain stem and olfactory bulbs, were dissected into cerebellum and non-cerebellum regions for microglia isolation. Single cell suspensions were prepared from brain tissues by mechanical dissociation using mesh of decreasing sizes from 250 μm to 70 μm and enriched for microglia by density gradient separation^72^. Briefly, the cell pellet was resuspended in 70 % (v/v) isotonic Percoll (1x PBS + 90 % (v/v) Percoll), overlayed with 37 % (v/v) isotonic Percoll and centrifuged with slow acceleration and no brake at 2000 xg for 20 min at 4 °C. The microglia-enriched cell population isolated from the 37 %-70 % interphase was diluted 1:5 in ice-cold PBS and recovered by cold centrifugation at maximum speed for 1 min in microcentrifuge tubes. The cell pellet was then stained with antibodies to microglial cell surface markers (CD11b-BV650, 1:200 Biolegend, #141723; CD45-BV786, 1:200, BD Biosciences #564225; CX3CR1-FITC, 1:100, Biolegend, #149019; CD11a, 1:20, BD Biosciences, #558191, TREM2-APC, 1:10, R&D Systems, #FAB17291N; CD33-PE, 1:20, eBioscience, #12-0331-82; CD115-BV711, 1:40, Biolegend, #135515) for isolation using the FACSAria™ III cell sorter. Microglia were defined as live/propidium iodide (PI)^−^ (Sigma-Aldrich, St. Louis, MO, #P4864), CD11b^+^, CD45^lo^, CX3CR1^+^ single cells and were negative for CD11a (Extended data Fig. 1). The XO4^+^ population gate was set using Methoxy-XO4-injected wild-type animals. X04^+^ and X04^−^ microglial populations were sorted separately for further analysis by bulk RNA-seq, nano LC-SWATH-MS (20,000 cells per sample) and single cell RNA-seq.

### viSNE Analysis

The Cytobank platform (Fluidigm, South San Francisco, California) was utilized to generate viSNE plots^35^ from Flow Cytometry Standard files. Analyses were performed on live/propidium iodide (PI)^−^ single cell population. A total of 40,000 events were sampled to generate viSNE maps. Seven fluorescent channels (CD11b, CX3CR1, CD45, CD115, CD33, TREM2 and Methoxy-X04) were engaged for dimensionality reduction. The run was performed 5 times to ensure the stability of the presented outcome.

### RNA-seq Library construction and sequencing

RNA extraction from 1-10 x10^4^ FACS-sorted microglia or iPS-derived microglia-like cells was performed on the QIAcube (Qiagen) using the RNeasy Micro Kit (Qiagen, #74004) and RNA quality was assessed using the Bioanalyser (Agilent RNA 6000 Pico kit; #5067-1513). The libraries were prepared using 0.5-2 ng of non-cerebellum microglia RNA samples with RIN value ≥ 7 and cerebellar microglia with RIN value ≥ 6. An 8 bp sample index (Supplementary table 8) and a 10 bp unique molecular identifier (UMI) were added during initial poly(A) priming and pooled samples were amplified using a template switching oligonucleotide. The Illumina P5 (5’ AAT GAT ACG GCG ACC ACC GA 3’) and P7 (5’ CAA GCA GAA GAC GGC ATA CGA GAT 3’) sequences were added by PCR and Nextera transposase, respectively. The library was designed so that the forward read (R1) utilizes a custom primer (5’ GCC TGT CCG CGG AAG CAG TGG TAT CAA CGC AGA GTA C 3’) to sequence directly into the index and then the 10 bp UMI. The reverse read (R2) uses the standard R2 primer to sequence the cDNA in the sense direction for transcript identification. Sequencing was performed on the NextSeq550 (Illumina), using the V2 High output kit (Illumina, #TG-160-2005) in accordance with the Illumina Protocol 15046563 v02, generating 2 reads per cluster composed of a 19 bp R1 and a 72 bp R2.

### Demultiplexing and Mapping

Sequencing reads were sample demultiplexed with Je demultiplex from the JE suite^73^ using sequence barcodes in Supplementary table 8. Short sequence unique molecular identifiers (UMIs) from read pair 1 of the demultiplexed sample sequencing reads were discarded from both sequencing read pairs with Prinseq (minimum length 9)^74^. Remaining UMIs were clipped with Je clip and added to the sequencing read header to allow UMI deduplication post read mapping. Demultiplexed UMI tagged sequencing reads were filter-trimmed with Trimmomatic^75^ and aligned to the mouse genome (GENCODE’s GRCm38 primary assembly annotation version vM15) using STAR^76^ (only sequencing reads from pair 2 were used for transcript quantification). Read deduplication based on UMIs was performed with Je markdupes and transcript read counts calculated with featureCounts^77^.

### Bulk RNA-seq analysis

The log2-transformed normalized gene expression from bulk RNA-seq was obtained using the Variance Stabilizing Transformation (VST) from the “DESeq2” package in R^78^. PCA (Fig. 1e) and hierarchical clustering (Extended data Fig. 2a) were then performed on the VST counts. To investigate if the sequencing batch had an effect on the gene expression, we performed a covariate analysis. For each covariate of interest (XO4, batch, region, age, genotype and gender), a likelihood ratio test (LRT) was performed using the “DESeq2” package, comparing the full model comprising all covariates and the reduced model which omits the covariate of interest. Thus, genes that are statistically significant under the LRT are genes whose variation in expression levels could be explained by the covariate of interest. The covariate analysis (Extended data Fig. 2b) revealed a large number of genes associated with batch (1020 genes, FDR < 0.01) and these genes significantly overlap with the region-related genes (*p*=3.0×10^−46^ by hypergeometric test) and XO4-related genes (*p*=2.2×10^−214^ by hypergeometric test, Extended data Fig. 2c, d). Thus, the batch covariate was included in subsequent analysis to account for batch effects. We also found that only 8 genes contribute to gender-related variation (FDR < 0.01). Thus, all subsequent analysis was performed excluding the gender covariate and both male and female microglial transcriptomes were analyzed together. The covariate analysis was then performed again without the gender covariate to identify genes that are specific to the XO4 covariate (Fig. 1h_i_). GO and KEGG term overrepresentation analysis were performed using the “gProfileR” package in R^79^. To generate the “gene cytometry” plots (Fig. 1f, g), a generalized linear model was constructed with the covariates XO4, batch, region, age and genotype. Separate pairwise differential expression analyses were then performed between XO4^+^ vs. XO4^−^, 4m vs. 1m and 6m vs. 1m microglia samples respectively. For each differential expression analysis and each gene, a gene score was then calculated as the product of the log2 fold change and negative log-transformed FDR, *LFC*\*-*FDR*, combining the effect size and statistical significance of the differential expression^80^. The gene scores for XO4 and age differential expression were then plotted to give the “gene cytometry” plots.

### Single cell sequencing

5,000 microglia from each population (including XO4^+^ and XO4^−^ microglia) were sorted into DMEM/F12 media (supplemented with 5% (v/v) FBS, 50 U/ml Penicillin and 50 µg/ml Streptomycin), centrifuged at 12,000 xg for 2 min at 4°C and resuspended in 35μl of PBS containing 0.04% (w/v) BSA (0.22 µm filtered). The samples were next diluted with nuclease-free water in accordance to 10X single cell protocol guidelines to achieve a target cell recovery of approximately 800 cells/sample. Single cell capture, RNA-seq library construction and sequencing were carried out at Micromon, Monash University using the 10X Genomics Chromium system (10X Genomics). Library construction was performed by poly-A selection from total RNA using 10X Chromium controller with Chromium Single Cell 3’ Reagent Kit V2 (10X Genomics, #PN-120237). Sequencing was performed on one High-Output lane of an Illumina NextSeq 550 (Illumina, California, USA) in paired-read 150 bp format. Chromium barcodes were used for demultiplexing and FASTQ files were generated using the Cellranger mkfastq pipeline. Alignment, filtering and UMI counting were performed using cellranger count. To improve detection of microglia, due to their low RNA content, cellranger reanalyse was used with the--force-cells option set at the inflection point when number of barcodes is plotted against the number of UMIs. Cells were manually filtered such that barcodes containing at least 10 counts corresponding to *Cx3cr1, P2ry12* or *Fcrls* genes were classified as microglia, resulting in a total of 991 cells from the 4 FACS-sorted microglial populations.

### Single Cell Analysis

The original mapped matrix dimensions were 10,484 genes by 991 cells. For quality control, various filtering steps were implemented. Genes without any counts in any of the cells were discarded. Cells were filtered by total counts and total features (genes) such that cells or genes below and equal to the 5th percentile were discarded. Next, cells with more than 10 % of their gene expression assigned to mitochondrial genes were discarded as these cells are likely to be undergoing apoptosis. Five sex-associated differentially expressed genes (*Xist, Ddx3y, Eif2s3y, Hsp90ab1, P4ha1*) identified in the bulk-RNA analysis that overlap with differentially expressed genes detected between 24m WT (female) and 6m WT (male) in our single-cell data were also filtered out. Cells not in G1 phase were also removed using scores calculated from cyclone^81^. Lastly, genes containing less than 1 count in at least 2 cells were removed from analysis, resulting in a dataset consisting of 6,685 genes by 893 cells. SCATER(version 1.6.1)^82^, SCRAN(version 1.6.6)^83^, and SINGLE CELL EXPERIMENT (1.0.0) were used for plotting PCAs and quality control plots^82^. Normalization was done by calculating Log_2_ Counts Per Million (CPM). Violin plots of DAM1 and DAM2 genes were obtained using Seurat’s VlnPlot function in R^84^.

### Feature Selection

For optimization, each feature selection method (M3DROP(version 3.5.0)^85^, Highly Variable Genes, Correlation-based, PCA-based, Depth Adjusted Negative Binomial (DANB)) was implemented before running SC3 (Single cell consensus clustering)(version 1.7.6)^40^. Rand index was calculated using MCLUST ^86^ to quantify accuracy of the feature selection method. DANB was found to have the highest rand index of approximately 90%. The number of feature genes was ascertained by calculating rand indexes after running SC3. We found that rand index does not increase significantly beyond the 25th percentile of genes used. Therefore, we used the top 1,671 (25th percentile) of the genes as our set of feature genes, which were optimal for discriminating subpopulations of cell in our dataset.

### Clustering

Clustering was performed using the single-cell consensus clustering (SC3) method^40^, which is based on unsupervised clustering of scRNA-seq data. The optimal number of clusters (k) was investigated using the *Sc3_estimate_k* function of SC3, and subsequently we set k = 4, achieving a rand index of approximately 91.5%. The Kruskal Wallis Test within SC3 (get_de_genes) was also used to detect differentially expressed genes across all 4 *a priori* labels and clusters (Plotly Technologies Inc. Collaborative data science. Montréal, QC, 2015. https://plot.ly).

### Differential expression and gene regulatory network analyses

#### Differential expression analysis

We utilized EdgeR^87^ via the EdgeRQLF function for pairwise differential expression analysis across two cell populations (i.e., between 2 *a priori* labels or 2 SC3 derived clusters), and size factors were calculated using computeSumFactors() from SCRAN. Multiple testing correction was implemented using the Benjamini & Hochberg (BH) correction and significant differentially expressed genes were called at the BH-adjusted *p*-value < 5% threshold.

#### Regulon identification

Gene regulatory network analysis was performed using single-cell regulatory network inference and clustering (SCENIC) method(version 0.1.7)^50^. SCENIC integrates a random forest classifier (GENIE3)(version 1.0.0) to identify potential transcription factor targets based on their co-expression with RcisTarget(version 0.99.0) for *cis*-regulatory motif enrichment analysis in the promoter of target genes (± 500 bp of the transcription start site (TSS)) and identify the regulon, which consists of a TF and its co-expressed target genes. The mus musculus 9 (mm9) motif database provided by the SCENIC authors were used. Finally, for each regulon SCENIC uses the AUCell (version 0.99.5) algorithm to score the regulon activity in each cell. The input for SCENIC was the 6,685 (genes) by 893 (cells) matrix obtained after filtering, as detailed above and gene expression is reported in Log_2_ CPM units. Unlike in the original SCENIC pipeline, we did not implement the 2-step filtering as suggested because the input matrix was already filtered using our own criteria. Otherwise, all parameters used for running were specified in the original SCENIC pipeline. The regulon activity matrix was binarized (giving 1/0 activity score for each cell) and the heatmap of the hierarchical clustering of the binarized matrix was plotted upon removing transcription factors with fewer than 100 genes (as these identified regulons are sporadically expressed in the binary heat map, and not clearly separated compared to larger regulons). Fig. 5i shows the top 2 regulons for XO4^+^ microglia and non-XO4+ microglia. In addition, we focused only on regulons that are active in more than 10 % of the cells. The identified regulons were visualized using *igraph* and *qgraph* R packages^88^.

#### Regulon annotation

The genes defining the regulon were input in STRING database(version 10.5)^89^ (www.string-db.org) and KEGG enrichment was carried out using the STRING web interface (background = all protein coding genes) (4^th^ June 2018). In order to obtain the relevant string protein-protein interactions (PPI), we downloaded the mouse string database and filtered for edges with combined score > 500 and experimental score > 0, giving us high confidence edges supported by experimental data. Text-mining setting was disabled for all analysis in order to minimize spurious gene associations.

### Pseudotime analysis

We used Diffusion Map in the destiny R package (version 2.6.1) for the pseudotime analysis^90^. Specifically, for the phagocytosing pseudotime, we used the list of 536 differentially expressed genes between 6m 5xFAD XO4^−^ and 6m 5xFAD XO4^+^ cells (FDR < 0.05). For ageing, we used 104 differentially expressed genes between 6m WT and 24m WT cells (FDR < 0.05). In order to plot the pseudotime, phagocytosis-specific and ageing-specific genes were defined as the non-overlapping genes between phagocytosis and ageing. This resulted in 474 phagocytosis genes and 42 ageing genes for diffusion map calculation. For defining pseudotime order, cells were ranked based on the first component of the diffusion map. For Extended Figure 5, we plotted the top 20 ageing-specific and top 20 phagocytosis-specific genes (based on absolute LogFC) ordered by their respective pseudotime.

### Projection Analysis

In order to determine the relation between our bulk and single-cell RNA-seq data, we projected our single cell data onto the bulk using Reference Component Analysis (RCA)^91^. Input units were in log_2_ (CPM) value, and no additional normalization or transformation was performed. Briefly, the expression profile of each single cell was projected onto each sample in the bulk RNA-seq data by calculating the Pearson correlation coefficient between the log_2_ (CPM) vector from scRNA–seq and bulk RNA-seq. For each cell, the Pearson correlation coefficients were *z*-score transformed and grouped by hierarchical clustering of the bulk RNA-seq data. The results of the projection analysis are reported in Extended data Fig 4b.

### Ingenuity Pathway Analysis (IPA)

To find upstream regulators of the regulons identified by SCENIC, we implemented Ingenuity Pathway Analysis (IPA)^51^ (27^th^ June 2018). For the *Hif1a* regulon (*n*=1,122 genes), we further pruned the regulon size as follows: first, we overlapped the genes in the regulon with the 1,671 feature genes resulting in common set of 371 genes; second, we derived the differentially expressed genes from the set using the Kruskal Wallis Test across all 4 *a priori* clusters (adjusted *p*-value = 0.05), which resulted in 203 genes. We used the set of 203 genes as input for IPA (five genes: *Gltscr2, Wbp5, Amica1, Myeov2*, and *0610011F06Rik* were not present in the IPA database) and their respective fold changes (FCs); here, we used the log_2_ FCs derived from the 5xFAD XO4^+^ vs 5xFAD XO4^−^ comparison. Next, we used IPA to predict the upstream regulators of the *Hif1a* regulon. As a first step, we extracted the regulators from the top five regulator effects networks robustly inferred by IPA (consistency score > 10) from the *Hif1a* regulon gene set. In doing this, we required the *Hif1a* gene to be included in the set, a direct downstream target of the regulated network. All upstream regulators must also have an absolute activation z-score higher than 2. We also required the regulated network to have a significant overlap with the *Hif1a* regulon gene set (*p*<0.05). The top ten upstream activators are reported in (Fig. 6a) and are ranked by their potential activation (z-score); the complete list of predicted activators presented in Supplementary Table 7. The cartoon diagram of the hypothesis generated by IPA was drawn using BioRender (https://app.biorender.com).

### Single nuclei sequencing analysis

#### Mapping single nuclei reads to the genome

Using the Grch38 (1.2.0) reference from 10x Genomics (see https://support.10xgenomics.com/single-cell-gene-expression/software/pipelines/latest/advanced/references for a detailed step-by-step description of the pipeline), we made a pre-mrna reference according to the steps detailed by 10x Genomics (https://support.10xgenomics.com/single-cell-gene-expression/software/pipelines/latest/advanced/references). Cellranger *count* was used to obtain raw counts. In all, our single nuclei data consists of 8 10x runs consisting of 4 AD runs and 4 control runs. Each run has 2 patients (see patient information above). 2 runs were discarded because of high neuronal enrichment (see Supplementary Table 9) possibly indicating neuronal contamination or technical artifacts.

#### Quality Control for expression matrix

The raw expression matrix was composed of 33 694 genes and 14 876 cells. Genes without any counts in any cells were filtered out. A gene was defined as detected if 2 or more transcripts were present in at least 10 cells. 100 PMI-associated genes, as defined by Zhu *et al* in the cerebral cortex, were removed^92^. For cell filtering, cells outside the 5th and 95th percentile with respect to number of genes detected and number of unique molecular identifier (nUMI) were discarded. In addition, cells with more than 10% of their genes being mitochondrial genes were filtered out. In addition, the matrix was normalized with scale factor of 10000 as recommended by the *Seurat* pipeline before *FindVariableGenes* was used to define variable genes with the parameters x.low.cutoff = 0.0125, x.high.cutoff = 3, and y.cutoff = 0.5. *ScaleData* was used to center the gene expression. Overall, the resulting filtered matrix consists of 10 850 genes and 13 214 cells.

#### Cell Type Identification

BRETIGEA^93^ is a R package which utilizes set of brain cell-type markers curated from independent human and mouse single cell RNA datasets for cell type proportion estimation in bulk RNA datasets. It can also be employed in single cell RNA datasets for the identification of cell types. For the reference datasets, BRETIGEA uses well-annotated and well-referenced datasets from ^34^ and ^94^. The 6 main cell type lineages identified by BRETIGEA are neurons, astrocytes, oligodendrocytes, microglia, oligodendrocyte progenitor cells (OPCs), and endothelial cells. We obtained markers from BRETIGEA for all 6 cell types and calculated a module score for each cell type using Seurat’s AddModuleScore function. This function calculates the average expression levels of each cell type subtracted by the aggregated expression of a control gene set of 100 randomly selected genes. Cell type identification was done in 2 steps.

Firstly, each cell was assigned a cell type based on the highest cell type score across all 6 cell types. Furthermore, we defined a cell as a hybrid cell if the difference between the first and second highest cell type scores are within 20% of the highest cell type score:

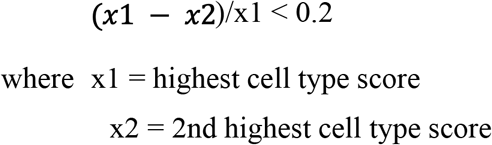

Secondly, for each cell type, we assumed normality of the gene score distribution and applied z-score transformation. Subsequently, in order to consider poorly identified cells i.e. cells with low cell type scores, for each cell type, cells with low cell type score (5th percentile and below) were relabeled as “unidentified” cells.

This robust statistical step identified 449 microglia.

#### Single human nuclei analysis

Seurat was used for normalization, scaling, and finding variable genes in the same sequence described in *“Quality control for expression matrix”*. Uniform manifold approximation and projection (UMAP) was recently recommended as a single cell data visualization tool ^95^. UMAP was calculated using RunUMAP command on the first 15 principle components. Differential expression between AD and control groups was then performed using the empirical Bayes quasi-likelihood F-tests (QLF) in the edgeR package (v3.20.8). Genes are deemed significantly differentially expressed if they have an FDR < 0.05. Out of the 40 human orthologs derived from taking the overlap between genes upregulated in AD microglia vs control microglia (341 genes, FDR < 0.05) and the XO4^+^ signature in mice (536 genes, FDR < 0.05), 30 of them (75.0%) have concordance of directionality. In contrast, out of the 1051 human orthologs derived from taking the overlap between genes upregulated in AD microglia vs control microglia (10537 genes, FDR <1.0) and the XO4^+^ feature genes in mice (1671 genes, FDR < 1), only 517 (49.2%) have concordance of directionality. Hypergeometric test, phyper(), was subsequently used to test for significance of concordance in directionality between the XO4^+^ signature in mice and the genes upregulated in AD human microglia. The formula used is phyper(30-1,517,1051-517,40, lower.tail = F). pheatmap was used to plot normalized gene expression of the 40 overlapping differentially expressed genes.

For pseudotime analysis, we defined phagocytosis and ageing genes as the top 10% of the differentially expressed genes for phagocytosis (XO4^−^ vs XO4^+^) and ageing (6m vs 24m), as previously, resulting in 167 ageing genes and 167 phagocytosis genes. For phagocytosis-specific and ageing-specific pseudotime, the overlap of 39 genes between the two signatures was removed. respectively. Seurat’s RunDiffusion was used for diffusion map calculation. bioMart^96^ (23rd January 2019) was used to convert mouse gene symbols to human orthologues. Overall, 128 ageing-specific and 128 phagocytosis-specific genes were used for pseudotime analysis. Pseudotime order was defined as the ranking of the first diffusion component. Similarly, phyper was used to calculate the respective p-values of the proportion of AD/Control cells lying beyond 50th percentile of the pseudotime. For example, the phyper formula for AD cells in the phagocytosis-specific trajectory is phyper(109-1,225,449-225,172, lower.tail = F).

#### Comparison with gene expression perturbed by small molecules

We consider the DEGs upregulated by TLR1-agonist administration but downregulated by rapamycin. We compared the log fold changes of *LGALS3, LY86, CTSB, HIF1A, LDLR, CD63, CTSL, CTSA* and *GAS7* in this dataset with those in XO4^−^ vs 6M WT and XO4^+^ 6M WT DE analysis. In addition, for supplementary figure 10e, we compared 866 DEGS upregulated by TLR1-agonist Pam3csk and downregulated by rapamycin with 1092 DEGS from the bulk RNA DE analysis between XO4^+^ and XO4^−^ genes (fdr <0.05, logFC>1). We subsequently plotted the overlap (64 genes) between these 2 sets of analysis to represent genes which are differentially expressed in both XO4^+^ microglia (vs XO4^−^ microglia) and upregulated by Pam3csk administration but downregulated by rapamycin. To calculate the p-value, we use a hypergeometric test where the population is taken to be the 4319 DEGS from the bulk RNA DE analysis between XO4^+^ and XO4^−^ genes (logFC>1). This set of genes has 92 genes overlapping with the 866 DEGS upregulated by Pam3csk and downregulated by rapamycin. The sample taken for the hypergeometric test is the 1092 DEGs which has 64 genes overlapping with the 866 DEGS upregulated by Pam3csk and downregulated by rapamycin. Overall, phyper formula used is phyper(64,92,4319-92,1092,lower.tail = F). bioMart was used to convert all mice genes to their human orthologues.

### In-solution tryptic digestion of proteins

Synaptosomes were prepared using the Procedure for Synaptic Protein Extraction from Neuronal Tissue” and Syn-PER Synaptic Protein Extraction Reagent (Thermo Fisher, #87793, containing a half tablet of cOmplete™ Protease Inhibitor Cocktail (Roche, #CO-RO) per 12.5 ml Syn-PER reagent) from 250 mg of frozen mouse brain tissue. Cell pellets were lysed (60 µL per 2×10^4^ microglial cells per sample; 150 µL per ~4 x10^5^ microglial cells for library preparation; 500 µL for bulk synaptosomes) in 1% (w/v) sodium deoxycholate (SDC, Merck) in 100 mM Tris pH 8.1, then boiled at 95 °C for 5 min. After centrifugation, sample supernatant was subjected to alkylation by addition of 40 mM chloracetamide (CAA, Merck) and incubation for 20 min at room temperature in the dark. Samples were digested by addition of porcine trypsin (enzyme to protein ratio of 1:100; for 2 x10^4^ microglial samples, 300 ng was used; Merck) and incubated overnight at 37 °C with shaking. The following day, the reaction was stopped and SDC precipitated through addition of formic acid to a final concentration of 1 % (v/v). Peptides were extracted through adding an equal volume of 100 % (v/v) water-saturated ethyl acetate, vortexing and centrifugation at maximum speed in a benchtop microfuge and transferring the aqueous phase to a new tube, with this entire step repeated once. Samples were then vacuum concentrated (Labconco Centrivap) and peptides purified by C_18_ ZipTips® (Merck) prior to analysis by LC-MS/MS or LC-SWATH-MS. Note that all samples were supplemented with 200 fmoles of each iRT peptide^97^ (Biognosys, #Ki-3002-1). For bulk synaptosome samples, peptides were further fractionated prior to LC-MS by reversed-phase HPLC using an Ettan ÄKTA micro HPLC system (GE Healthcare), as described elsewhere^98^.

### LC-MS/MS and spectral library generation

Spectral libraries were generated from mass spectrometry of tryptic peptides derived from a combination of microglia cells and synaptosomes. Purified peptides were analysed on a TripleTOF® 6600 mass spectrometer (SCIEX) equipped with an on-line Eksigent Ekspert nanoLC 415 (SCIEX). Following autosampler injection, samples were subjected to trap-elution through loading onto a trap column (Eksigent nanoLC trap, #5016752; ChromXP C18, 3 µm 120 Å, 350 µm x 0.5 mm [SCIEX]) at 2 µl/min for 10 min in loading buffer (2 % (v/v) acetonitrile in water supplemented with 0.1 % formic acid) followed by separation at 300 nl/min across an analytical column (Eksigent nano LC column, #805-0012; ChromXP C18, 3 µm 120 Å, 75 µm x 15 cm [SCIEX]) equilibrated in 98 % buffer A (0.1 % (v/v) formic acid in water) and 2 % (v/v) buffer B (80 % (v/v) acetonitrile in water supplemented with 0.1 % (v/v) formic acid) followed by increasing concentrations of buffer B. Specific gradient conditions were: increase from 2-10 % (v/v) buffer B from 0 to 2 mins, then 10-40 % (v/v) buffer B from 2 to 152 mins, then 40-50 % (v/v) buffer B from 152 to 154 mins, then 50-99 % (v/v) buffer B from 154 to 157 mins, then hold at 99 % (v/v) buffer B from 157 to 167 mins, then 99-2 % (v/v) buffer B from 167 to 168 mins, followed by re-equilibration at 2 % (v/v) buffer B until the end of the run. The mass spectrometer was operated in information dependent acquisition mode using the following settings: MS1 accumulation time of 200 msec, scan range of 300-1800 m/z; for MS2, a switch criteria was used of the top 20 precursors exceeding 40 counts with charge state from 2 to 5, rolling collision energy and with ions excluded for 30 sec after two occurrences; MS2 accumulation time was set to 150 msec and with a scan range of 80-2000 m/z. Acquired spectra were searched using ProteinPilot™ v5 (SCIEX) against the complete reference mouse proteome (Uniprot, 201707 build) and the resultant search was imported into Skyline v3.7.11317^99^.

### LC-SWATH-MS

For data-independent acquisition, purified peptides were analysed on a TripleTOF® 6600 mass spectrometer (SCIEX) using the same LC setup and conditions as above, with the exception that the mass spectrometer was operated in SWATH-MS^100^ mode, using the following conditions: initial MS1 scan across 400-1250 m/z with accumulation time of 150 msec, followed by 100 variable SWATH windows (calculated using the Variable Window Calculator Excel tool, downloaded from http://sciex.com/support/software-downloads) spanning a range of 400-1241 m/z with a 1 Da overlap and each with an accumulation time of 25 msec. Rolling collision energy was used, with a collision energy spread of 5.

### SWATH-MS data analysis

SWATH-MS data was analysed using Skyline v3.7.11317 against the generated microglial and synaptosome spectral library, applying predicted retention times from the included iRT peptides detected within each sample in order to aid peak picking. Peak scoring was then re-trained within Skyline following the addition of shuffled decoy peptides. Data were initially refined through accepting peptides with an absolute ppm <10 and, for each peptide, there needed to be at least one sample of the set with a peptide dotp of >0.8; peptides not meeting these criteria were excluded. Subsequently, all remaining peptides were subjected to manual interrogation followed by exporting integrated peak areas into Microsoft Excel 2016 for further processing. The list of peptides used for signal normalization is listed in Supplementary table 2. Peptides corresponding to 304 of the XO4^+^ covariate genes were above the limit of detection. Samples and proteins were clustered using one minus Spearman correlation, and data expressed as a heat map of log2-transformed normalized fold changes compared to WT microglia using GENE-E software v3.0.215 (Broad Institute).

### Organotypic hippocampal slice cultures

Organotypic brain slice cultures (OSHCs) were adapted from published protocols^101^. On day 0, brains from 6-month-old 5xFAD and WT animals were coronally sectioned through the hippocampus on a vibratome (Leica; settings: speed=0.4 mm/s and amplitude=1.00 mm) in ice-cold cutting medium (MEM 1X, Life Technologies) containing 10 mM Tris, 50 U/ml Penicillin and 50 µg/ml Streptomycin) to obtain 400 μm thick brain slices. Brain slices were cultured *in vitro* with culture medium (25% (v/v) MEM 2x, 25% (v/v) HBSS 1x (Life Technologies), 25% (v/v) Horse serum (Life Technologies) with 10 mM Tris, 25 U/ml Penicillin and 25 µg/ml Streptomycin and 0.455% (v/v) 7.5% NaHCO_3_ aqueous solution) on Millicell Cell Culture Insert (Merck) at air-medium interface. The media was completely replaced every second day. After 3 days of resting, Aβ plaques on brain slices were stained using an alternative fluorescent amyloid plaque labelling dye, NIAD-4 (10 μM, BioVision, #2710) for 3 h, prior to the addition of *ex vivo* microglia enriched fraction isolated from 6-month-old 5xFAD and WT animals. Microglia-rich fractions enriched by Percoll gradient (described in Section 4.2.2) were stained with CFSE (final concentration 5 μM; Life Technologies) for 20 min at 37 °C, and 2×10^4^ cells were added per hippocampus onto NIAD-4-stained hippocampal slice cultures for 5 days. As a control, synaptosome-labelled pHrodo-red particles were added to OHSCs for 5 days. Endogenous and replenished microglia were purified from OSHCs by mechanical dissociation using 70 μm mesh and enriched for microglia by density gradient centrifugation in 30 % (v/v) isotonic Percoll at 1000 xg for 15 min. Cell pellets were incubated with Fc block (1:200; BD Biosciences, #553141) for 15 min on ice prior to staining with CD11b-PE (1:50) and CD45-BV786 (1:200) for 15 min. Cells were washed once in PBS and resuspended in 400 μl Zombie IR dye (1:1000; Thermo Fisher Scientific) for live cell discrimination. Endogenous (CFSE^−^) and exogenous (CFSE^+^) microglia (single, live, CD11b^+^, CD45^lo^) that were either positive or negative for NIAD4 were sorted into 96 well plates (10 cells/well) containing 10 μl of Lysis Buffer from the Single Cell to Ct kit (Thermo Fisher Scientific), using the FACSAria™ III cell sorter. As a control, pHrodo-red-containing microglia were also sorted from the slices. cDNA synthesis and pre-amplification (18 cycles) were performed in accordance to manufacturer’s instructions and pre-amplified cDNA was diluted 5-fold prior to qPCR. The primers and probes used are listed in Supplementary table 10.

### Differentiation to iMGLs

#### iHPC Differentiation Base Medium

IMDM (50%; Thermo Fisher Scientific), F12 (50%), ITSG-X, 2% v/v, Thermo Fisher Scientific), L-ascorbic acid 2-Phosphate magnesium (64 μg/ml; Sigma), monothioglycerol (400 μM; Sigma), PVA (10 μg/ml; Sigma), Glutamax (1X; Thermo Fisher Scientific), chemically-defined lipid concentrate (1X; Thermo Fisher Scientific), non-essential amino acids (NEAA; 1X; Thermo Fisher Scientific), Penicillin/Streptomycin (P/S; 1% V/V; Thermo Fisher Scientific). Use 0.22 μm filter.

#### iMGL Differentiation Medium

phenol-free DMEM/F12 (1:1), ITS-G, 2%v/v, B27 (2% v/v), N2 (0.5%, v/v), monothioglycerol (200 μM), Glutamax (1X), NEAA (1X), and additional insulin (5 μg/ml; Sigma), filtered through a 0.22 μm filter; supplemented with M-CSF (25 ng/ml; Milteniy biotec), IL-34 (100 ng/ml; Milteniy biotec), and TGFβ-1 (50 ng/ml; Milteniy biotec) and cholesterol(1.5 μg/ml; Avanti Polar Lipids, ^102^).

The protocol for iMGL derivation was adapted from ^55^ with modifications from the StemDiff Hematopoietic Kit (Stem Cell Technologies, #05310). H9 CX3CR1-TdTomato cells were cultured on vitronectin (Thermo Fisher Scientific, #A14700)-coated T25 flask (Sarstedt, #83.3910.002) in E8 medium (Thermo Fisher Scientific, #A1517001). 2 days prior to differentiation, cells were detached in 0.5 mM EDTA and 40-80 colonies /well were seeded into a 12 well plate in E8 medium. On day 0, E8 medium was exchanged for 1 mL of supplemented iHPC Differentiation Base Medium per well. iHPC Differentiation Base Medium supplemented with FGF2 (50 ng/ml), BMP4 (50 ng/ml), Activin-A (12.5 ng/ml), ROCKi (1 μM) and LiCl (2 mM), and incubated in a hypoxic incubator. On day 2, medium was changed to 1 ml of iHPC Differentiation Base Medium supplemented with FGF2 (50 ng/ml) and VEGF (50 ng/ml) and incubated in a hypoxic incubator. On day 4, media was changed to 1 ml iHPC Differentiation Base media containing FGF2 (50 ng/ml), VEGF (50 ng/ml), TPO (50 ng/ml), SCF (10 ng/ml), IL-6 (50 ng/ml), and IL-3 (10 ng/ml) and incubated under normoxia. Half the medium was replaced on days 5 and 7. On day 10, the supernatant containing the HPCs was collected, centrifuged (300 x g for 5 min at room temperature), then 0.5ml cell-containing media was replaced and supplemented with 0.5ml fresh media. On day 12, the supernatant containing HPCs was collected and plated onto matrigel (1:100; hESC-qualified Matrix, LDEV-Free, Falcon, #354277)-coated 12 well plates at 4×10^4^ cells/well in iMGL complete differentiation medium. Every two days, each well was supplemented with 0.5 ml per well of complete differentiation medium, and at day 22 a 50% media change was performed. From day 35, iMGLs were cultured in complete iMGL media supplemented with CD200 (100 ng/ml; Novoprotein) and CX3CL1 (100 ng/ml; Peprotech) for an additional three days. iMGL experiments were performed at d22-23 of differentiation. iMGLs were stimulated with human BMP9 (20ng/ml; Biolegend), Pam3csk (100 ng/ml; Invivogen), Poly:IC (1000 ng/ml; Invivogen) and/or rapamycin (5 nM; Sigma) for 24h prior to RNA isolation or collection of culture supernatant.

### Immunofluorescence staining

Right brain hemispheres from Methoxy-X04 injected PBS-perfused mice were fixed in 4% PFA overnight, followed by immersion in 30% (w/v) sucrose solution for 48 h and frozen with liquid N_2_. Samples were stored at −80°C prior to sectioning. Frozen hemispheres were cryostat-sectioned into 60 or 20 μm thick sections onto slides for histological staining. Sections were blocked for 1.5 h in PBST (containing 2% (w/v) BSA and 0.5% (v/v) Triton-followed by 1 h incubation with 0.5% (v/v) Mouse on Mouse (M.O.M.™) Blocking Reagent (Vector Laboratories, #MKB-2213-1). Sections were then stained with primary antibodies, including the rabbit anti-Iba-1 (1:500, #019-19741, Wako, Virginia, USA), mouse anti-PSD95 (1:160, #MAB1596, Merck Millipore), mouse anti-6E10 (1:500, #803001, Biolegend), mouse anti-GAD65 (1:1000, #ab26113, Abcam) or rat anti-C3 (1:50, #ab11862, Abcam) overnight at RT followed by Alexa Fluor 488 goat anti-rabbit IgG (H+L) (1:250, #A11008, Life Technologies), Alexa Fluor 635 goat anti-mouse IgG (H+L) (1:250, #A31575, Life Technologies) or Alexa Fluor 568 donkey anti-mouse IgG (H+L) (1:500, #A10037, Life Technologies), respectively, for 2 h at RT. Sections were then mounted with Mowiol mounting medium. Paraffin embedded human brain sections were de-waxed for antigen retrieval with 90% formic acid (5 min) followed by citrate boiling (45min, 98°C DAKO Citrate Buffer, DAKO PT Link). Sections were then blocked using 0.1% (w/v) Sudan Black B in 70% ethanol (5 min) followed by PBST (containing 2% (w/v) BSA and 0.5% (v/v) Triton-X; 1 h). Sections were stained with primary antibodies, including the rabbit anti-Iba-1 (1:500, #019-19741, Wako), goat anti-PSD95 (1:50, # ab12093, Abcam) or mouse anti-6E10 (1:500, #803001, Biolegend) overnight at RT followed by Alexa Fluor 488 goat anti-rabbit IgG (H+L) (1:250, #A11008, Life Technologies), Alexa Fluor 647 donkey anti-goat IgG (H+L) (1:250, #A21447, Life Technologies) or Alexa Fluor 568 donkey anti-mouse IgG (H+L) (1:500, #A10037, Life Technologies), respectively, for 2 h at RT. Sections were then mounted with Mowiol mounting medium.

### Confocal imaging and image analysis

Mouse sections were imaged on a Leica SP8 confocal microscope using a 63x oil 1.4 NA objective and 2x zoom with 1024×1024 resolution, resulting in a pixel size of 90 nm. 30-40 μm z-stacks were acquired with a 0.3 μm z-step size, using sequential scans with 3x averaging at 488nm and 647nm wavelengths and 1x averaging at 405nm wavelength. For human brain sections, microglia were analysed from four 4.5-5 μm z-stacks per patient, obtained at 63x, 1x zoom, 2048×2048. For quantitation of PSD95 or GAD65 engulfed by microglia or C3 co-localization with microglia, 3D rendering of confocal images was performed using surfaces, spots and cell functions and batch analysis in Imaris (v8.3.1). XO4+ microglia were defined as IMARIS-rendered surfaces in the Iba1 channel with mean fluorescence intensity above threshold in the Methoxy-XO4 channel. Experimenters were blinded to the genotype during image acquisition and processing.

### qRT-PCR

qPCR was performed using the Biomark Fluidigm 96.96 protocol. Briefly, assays (Supplementary table 10) and samples were combined in a 96.96 Dynamic array IFC according to Fluidigm® 96.96 Real-Time PCR Workflow Quick Reference PN 6800088. 5 μl of each assay at a final concentration of 10x was added to each assay inlet port and 5 μl of diluted sample was added to each sample inlet port according to the ChipPipetting Map. The data was analysed with Fluidigm Real-Time PCR analysis software (V4.1.2). The limit of detection was set to 30. Samples with a Ct value for *Actb* (Mm00607939_s1) outside the 15-25 range were excluded from further analyses. log2 transformed ΔCt values for the 42 detected genes were used for clustering by SC3 and relationships between samples were visualized by SPRING (https://kleintools.hms.harvard.edu/tools/springViewer), which uses a k-nearest-neighbor graph rendered using a force directed layout^103^. Plots were generated using the ggplot2 function in R. For validation of *HIF1A* regulon in human ES-derived iMGLs, 400 ng RNA was reverse transcribed using SuperScript™ II Reverse Transcriptase (Thermo Fisher Scientific, #18064014). qPCR was performed using TaqMan assays (*SPP1*, Hs00959010_m1; *HIF1A*, Hs00153153_m1; *GAPDH*, Hs02758991_g1; *APOE*, Hs00171168_m1; *TREM2*, Hs00219132_m1; *CX3CR1*, Hs01922583_s1; housekeeping gene *SNRPD3*, Hs00188207_m1) in a Roche LightCycler® 480 (Roche).

### Cytometric Bead Array (CBA)

CBA was carried out using the BD (New Jersey, USA) CBA human flexi kit using a protocol modified from the manufacturer’s protocol. 5 μl of each standard (highest concentration at 5000 pg/ml in assay diluent) and sample were incubated with 5 μl of capture bead mix (containing 0.1 μl of each cytokine Capture Bead diluted in Capture Bead Diluent) for 1 h in a 96 well V-bottom assay plate. This was followed by the addition and incubation with detection reagent mix (containing 0.1 μl of each cytokine PE Reagent diluted in Detection Reagent Diluent for 1 h in the dark. Each well was then washed once with 200 μl of Wash Buffer, and beads were resuspended in 80 μl of Wash Buffer for analysis by FACS using the LSRFortessa X-20 (BD Biosciences). At least 200 single bead events from each cytokine population were collected. Results obtained were analysed using the FCAP Array Software Version 3.0 (BD).

### Human primary microglia culture and transfection

Primary human microglia isolated from cortex (Celprogen, #37089-01) were cultured on matrigel-coated 12 well plates at 5×10^5^ cells per well in iMGL differentiation media at 37 °C in 5 % CO_2_. Cells (up to passage 5) were transfected for 48 h with dox-inducible GFP-tagged Gateway-generated Piggybac expression constructs with inserted cDNA encoding human *HIF1A* and/or *ELF3* open reading frames using Glial Mag transfection kit (Oz Biosciences, #GL00500) according to manufacturer’s instructions. Primary microglia were co-transfected with plasmids encoding Hybase transposase and *rtta* at a ratio of 1:2:2 (Hybase: rtta: ORF).

### Preparation and treatment with fibrils

Aβ_1-42_ (Bachem, #4014447.5000) monomers were prepared by HFIP solubilization and aliquots were stored at −20 °C over desiccant prior to use. For preparation of aggregated fibrils, Aβ_1-42_ was freshly resuspended 5 mM in DMSO at RT and diluted to 100μM final Aβ with 10 mM HCl and 150 mM NaCl at RT made in 18MΩ sterile water. Following 15 sec vortex, Aβ was incubated at 37 °C for 24 hrs. 200 nM Aβ was added to primary microglia or iMGLs for 48, 96 or 144 h.

### Phagocytosis of pHrodo *E. coli* and synaptosomes

Synaptosomes were isolated from WT mouse brain tissue or human brain (obtained from Victorian Brain Bank), according to the Syn-PER Synaptic Protein Extraction Reagent (Thermo Fisher Scientific, #87793) protocol. The protein concentration was measured by nanodrop, and mouse synaptosomes were labelled with pHrodo™ Red succinimidyl ester (Thermo Fisher Scientific, #P36600) as described by ^104^. Human synaptosomes were labeled with blue fluorescent 2.0 µm FluoSpheres™ Carboxylate-Modified Microspheres (Life technologies, F8824) according to manufacturer instructions. Mouse pHrodo-conjugated synaptosomes were resuspended at 3.5 μg/μl in 5% DMSO in PBS and stored at −80 °C until use and human pHrodo-conjugated synaptosomes were resuspended at 5.5 mg/ml in 1 % (w/v) BSA and stored at 4 °C until use. To examine the phagocytic properties of the XO4^+^ and XO4^−^ microglia populations, the microglia-enriched cell suspension were isolated from Methoxy-XO4-injected mice as described above were seeded in 96 well plates. Following 30 min of resting at 37 °C and 5 % CO2, microglia were incubated with pHrodo-conjugated synaptosomes (4.25 μg per well) or pHrodo™ Green *E. coli* BioParticles™ Conjugate (Thermo Fisher Scientific, #P35361, 66.7 ng per well). iMGLs or primary human microglia were incubated with conjugated-synaptosomes (3.44 mg/ml). Cells were collected after 45 min incubation at 37 °C and 5 % CO_2_ and stained with antibodies to microglia cell surface markers (CD11b-PE-Cy7, 1:200, Biolegend, #101216; CD45-APC-Cy7, 1:200, Tonbo Biosciences, #25-0459-T100). XO4^+^ and XO4^−^ microglia uptake of pHrodo-conjugated synaptosomes or pHrodo™ Green E. coli BioParticles™ Conjugate was analysed using the FACSAria™ III cell sorter.

### Statistics

Differences between 2 groups were compared by two-tailed *t-*test, and more than 2 groups, by one-way ANOVA and Tukey’s or Sidak post hoc tests, as appropriate. To analyze significance of gene enrichment of sensome genes in the XO4^+^ signature, we used hypergeometric test. The hypothesis that plaque distal human microglia internalize more PSD95 than plaque-associated microglia was tested by paired one-tailed one-sample *t-*test.

### Data availability statement

Bulk RNA-seq and Single cell-seq and single nuclei-seq data from this study are available from: http://www.systems-genetics.net/assets/files/Grubman_et_al_2018_bulkRNA.zip

Proteomics raw data are included in Supplementary table 2. Raw sequencing reads have been submitted to the NCBI’s Sequence Read Archive (SRA, https://www.ncbi.nlm.nih.gov/sra).

## ACKNOWLEDGMENTS

The authors acknowledge Flowcore, Monash Micro Imaging, Monash Health Translation Precinct Medical Genomics Facility, Micromon, Monash Histology Platform and Australian Research Laboratories/Monash Animal Research Platform, Monash University, for the provision of instrumentation, training and technical support. We thank Dr. Steve Chai and Genevieve Buckley for their expertise with Imaris, Angela Vais for her advice regarding tissue sections, Dr. Steven Firth for provision of confocal imaging expertise. We thank Dr. Gurpreet Kaur (10X Genomics) for her insight regarding single cell sequencing analysis. We thank Kirill Tsyganov for a pipeline for mapping bulk RNA-seq data. The Australian Regenerative Medicine Institute is supported by grants from the State Government of Victoria and the Australian Government. A.G was funded by a NHMRC-ARC Dementia Fellowship and Dementia Australia Research Foundation Grant. J.M.P. was funded by a Sylvia-Charles Viertel Fellowship. J.M.P., C.P., J.H. and A.G. received a Monash University JMP Grant. Part of this work was funded by a Monash Network of Excellence grant. G.S. was funded by the Yulgilbar Foundation.

## AUTHOR CONTRIBUTIONS

AG and JMP conceived the study and designed experiments and together with EP designed the bioinformatics analysis. XYC, AG and GS performed microglia isolation, FACS, viSNE, immunostaining, OHSC and human microglia-like cell and primary microglia experiments with support from and ZA. CM performed pathological assessment of human control and AD cases. GC and EP performed single cell RNA-seq, pseudotime. IPA and regulon analyses. JFO and OJLR performed the bulk RNA-seq analyses. NC and AP performed proteomics experiments, SYC provided reagents and technical assistance. JP, RS, SB, DVL and RL worked up the protocol for single nuclei sequencing from human brain. FR mapped RNA-seq data and SW extracted single cell data. AG, XYC, GC, EP and JMP wrote the manuscript. All authors approved of and contributed to the final version of the manuscript.

