## Extended data Fig for "Mouse and human microglial phenotypes in Alzheimer’s disease are controlled by amyloid plaque phagocytosis through Hif1α"

### Extended data Fig 1

a

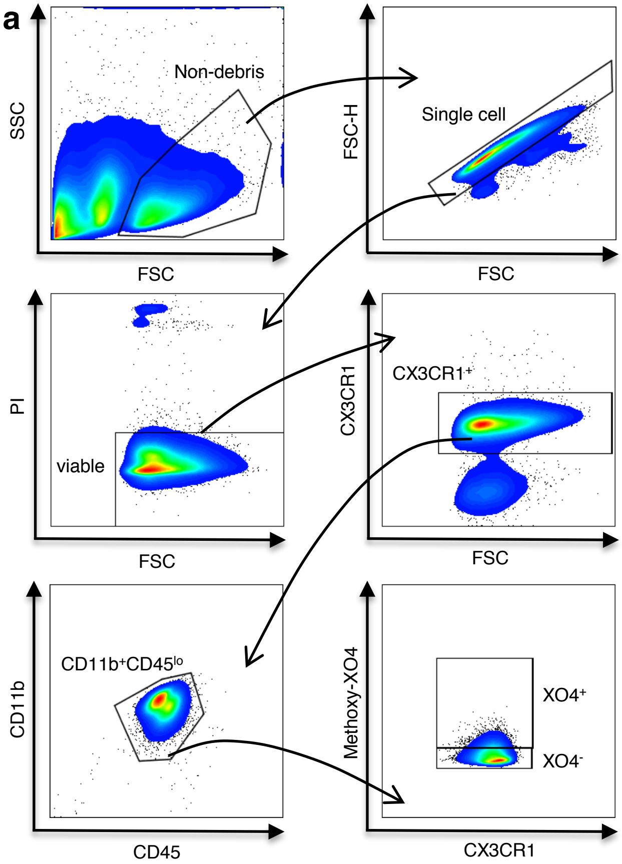


b

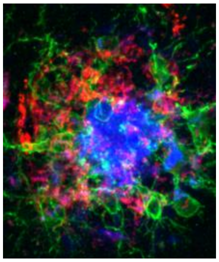

MeXO4/Iba1/6E10

**Extended data Fig. 1a** FACS gating strategy for sorting $XO4^+$ and $XO4^-$ microglia. **b**, Methoxy-XO4 labels the core of $6E10^+$ plaques, representative image from *n*=6 mice.

### Extended data Fig 2

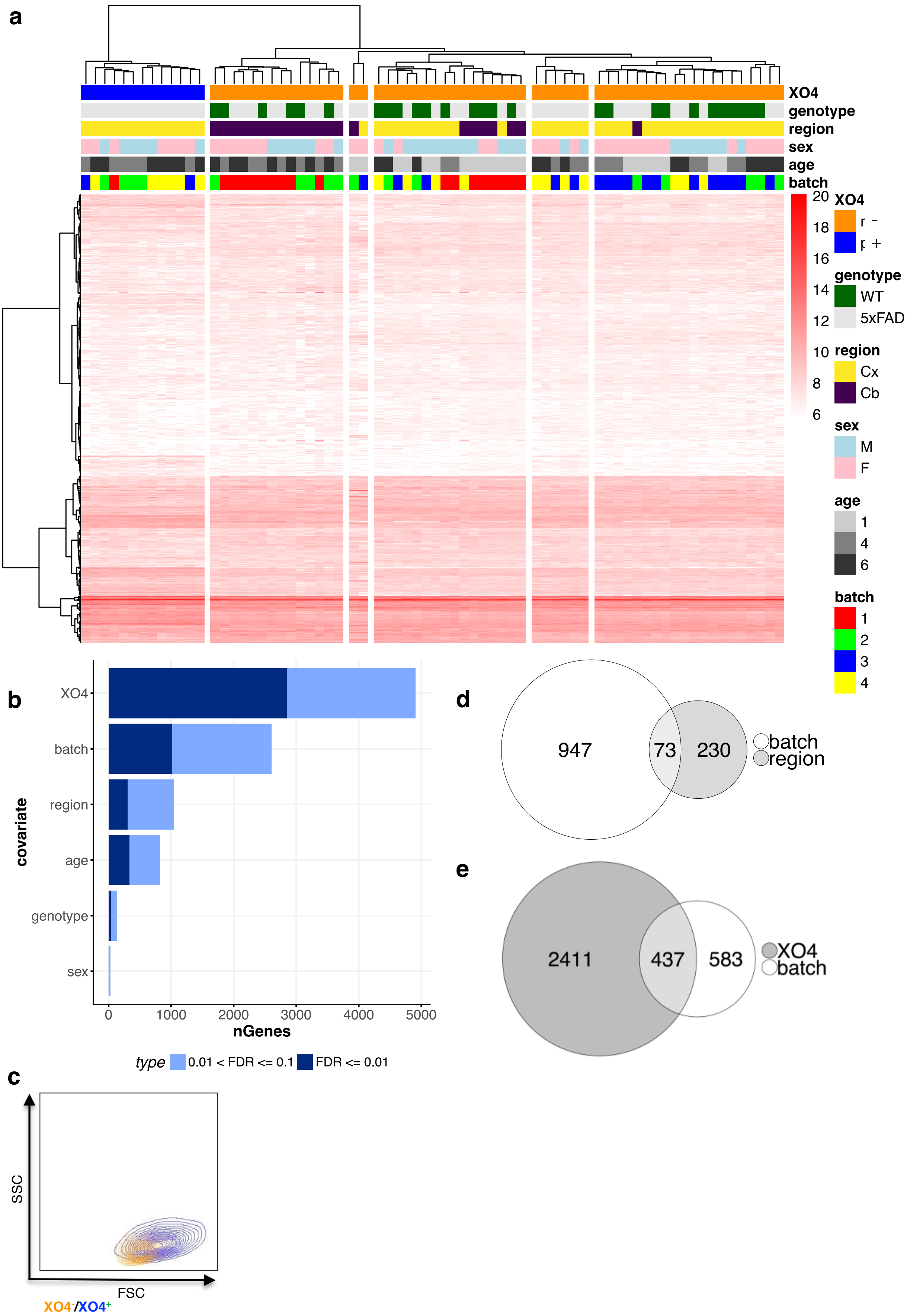

**Extended data Fig. 2a** Hierarchical clustering using ward.D2 linkage and Euclidean distance of bulk RNA-seq data. **b,** the number of genes for which expression levels could be explained by each covariate. **c**, FSC by SSC FACS plot showing increased size of $XO4^{+}$ microglia compared to $XO4^{-}$ microglia, representative of $n$=16 mice. **d-e,** Venn diagram showing the overlap between the batch and region (**d**) and batch and XO4 (**e**) covariate genes.

Extended data Fig 3

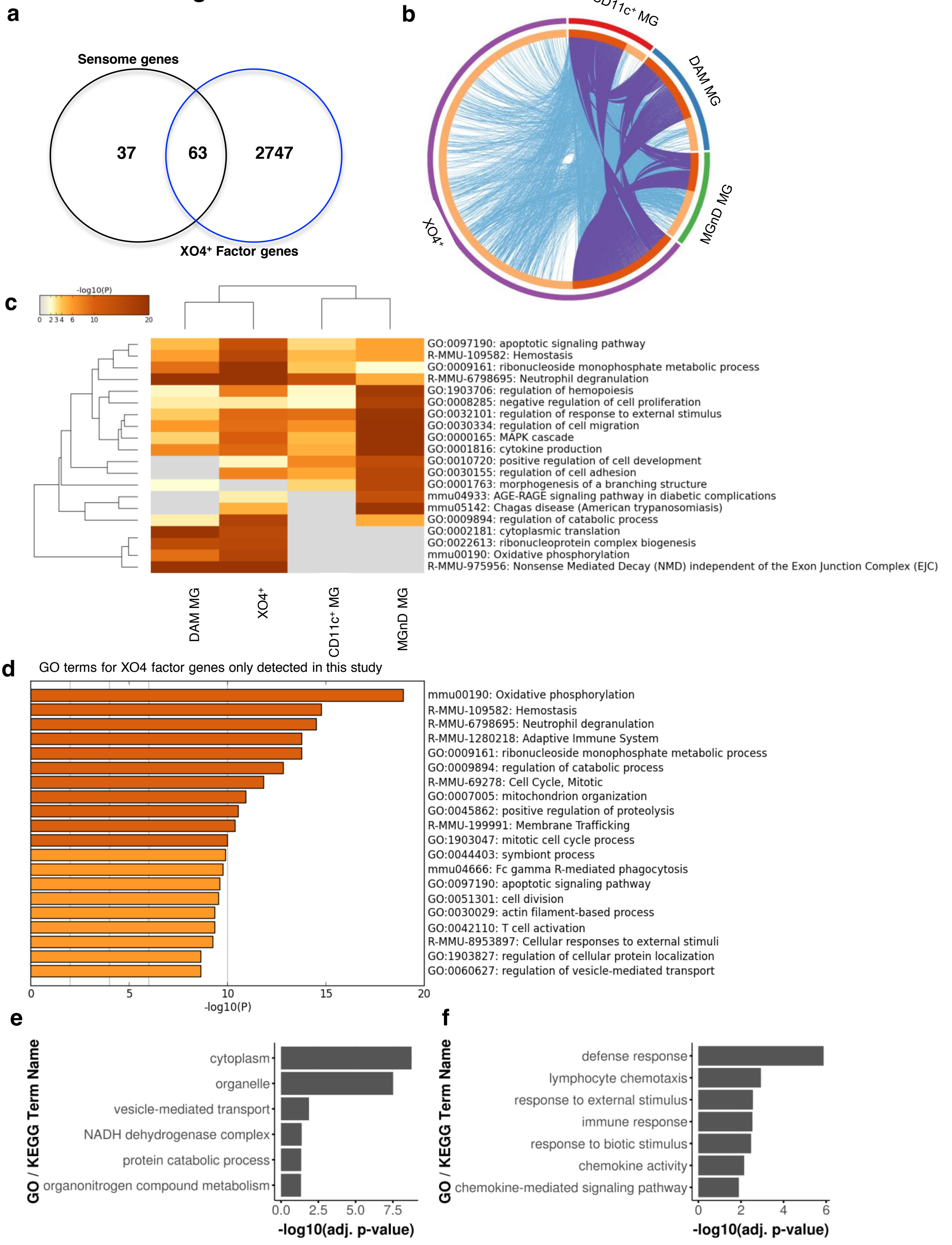

**Extended data Fig. 3a,** Overlap between microglial sensome genes and $XO4^+$ covariate genes, $p$=6.1x$10^{-11}$ by hypergeometric distribution. **b,** Circos diagram showing overlap in differentially expressed genes between WT microglia and $XO4^+$ microglia identified in this study by SPIA (bulk) compared to 10X single cell sequencing, and compared with DAM microglia[28], $CD11c^+$ microglia[29] and MGnD microglia[30]. Purple lines represent genes that are identified across multiple datasets, whereas blue lines represent gene ontologies that are identified across multiple datasets. **c**, Heatmap of enriched gene ontology terms in the microglial populations identified in each study, colored by *p*-values (legend). **d,** Heatmap of enriched gene ontology terms for $XO4^+$ covariate genes only identified in this study, colored by *p*-values (legend). Analysis was performed in Metascape. Gene ontology analyses were performed in Metascape. **e-f,** GO and KEGG terms associated with region (**e**) and age (**f**) covariate genes.

**Extended data Fig 4**

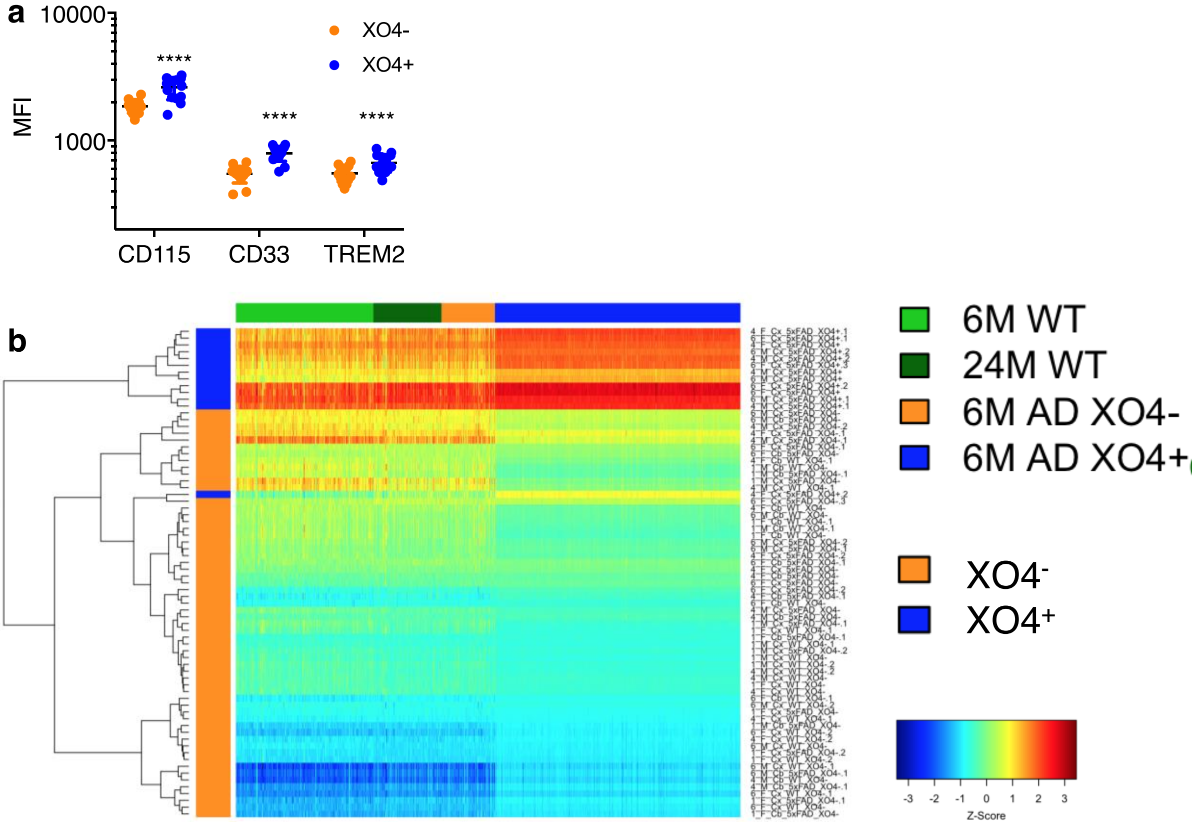

**Extended data Fig. 4a**, the median fluorescence intensity (MFI) of CD115, CD33 and TREM2 in 6m 5xFAD FACS-gated $XO4^-$ compared to $XO4^+$ microglia, pooled males and females, *n*=14 mice. **b,** Projection of single cell RNA-seq data onto bulk RNA-seq data using Reference Component Analysis (RCA).

#### Extended data Fig 5

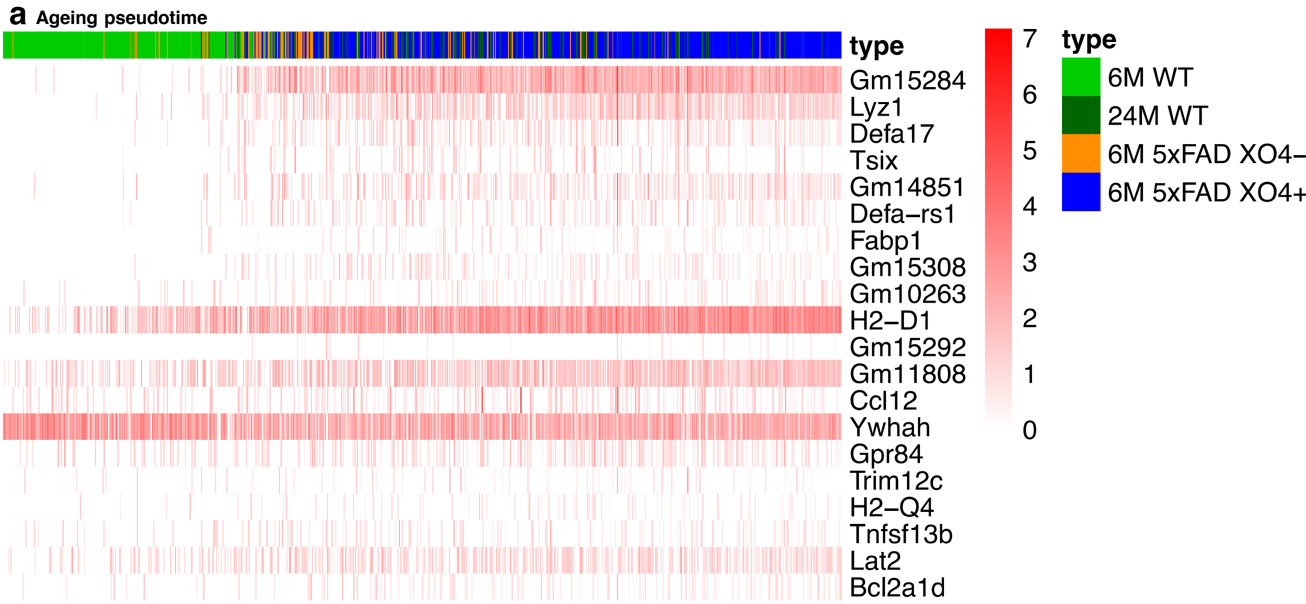


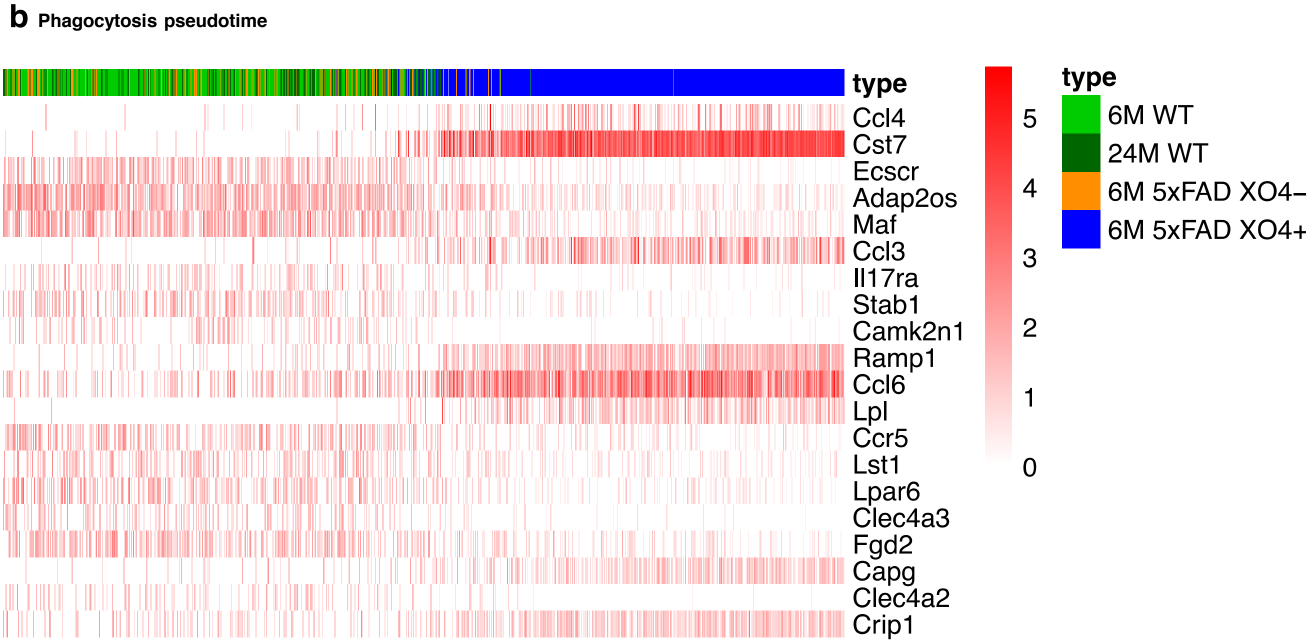

**Extended data Fig. 5**, $Log_2$CPM Expression Heatmap of top 20 trajectory-specific DE genes (24M WT vs 6M WT and $XO4^+$ vs $XO4^-$) ordered by **a**, age Pseudotime or **b,** phagocytosis pseudotime, respectively.

### Extended data Fig 6

**a** Stage 1 DAM genes

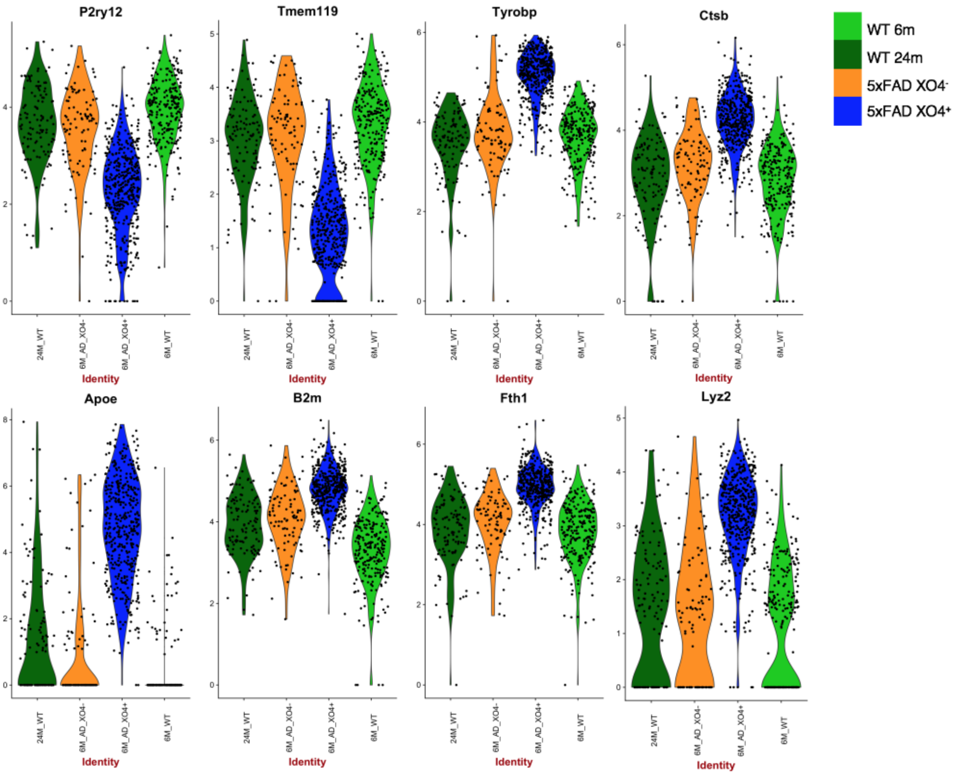


**b** Stage 2 DAM genes

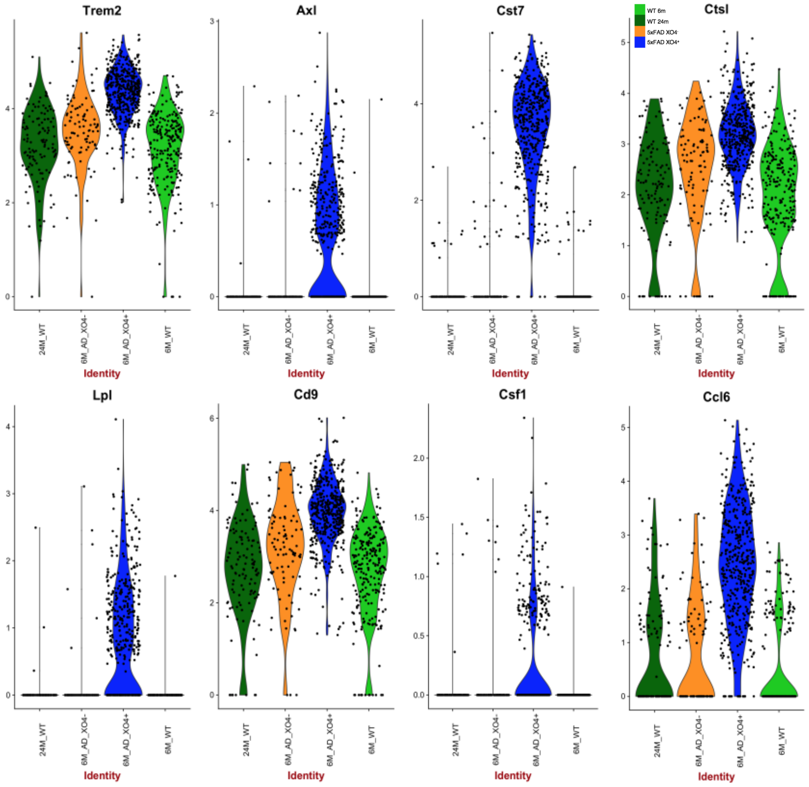

**Extended data Fig. 6**, Violin plots showing $Log_2$CPM expression of **a,** Stage I DAM and **b,** Stage II DAM genes[28] in *a priori* cell clusters.

### Extended data Fig 7

**a** Genes downregulated in NIAD4+

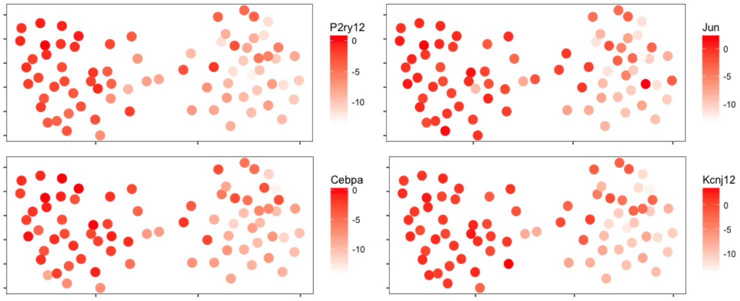


**b** Genes upregulated in NIAD4+

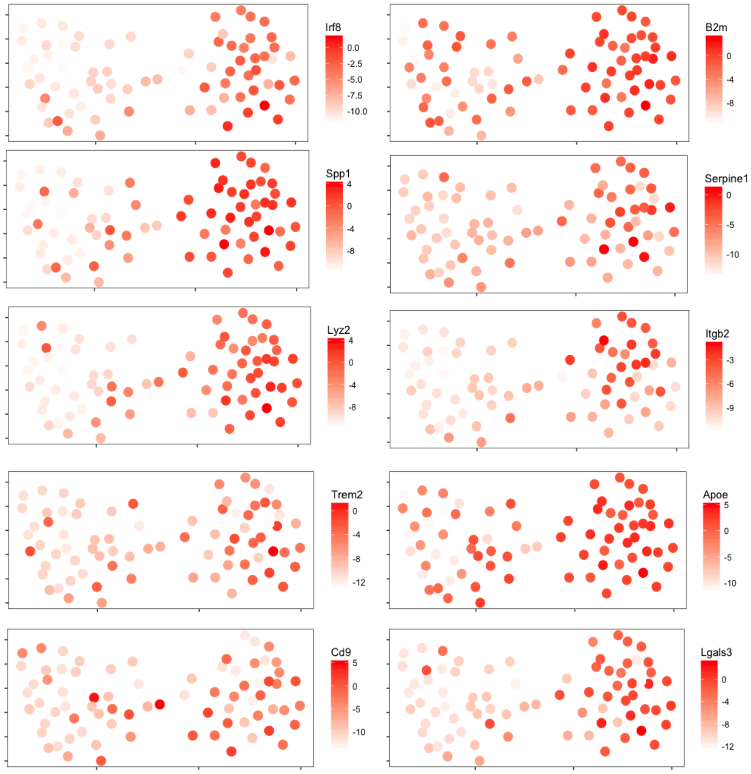


**c**

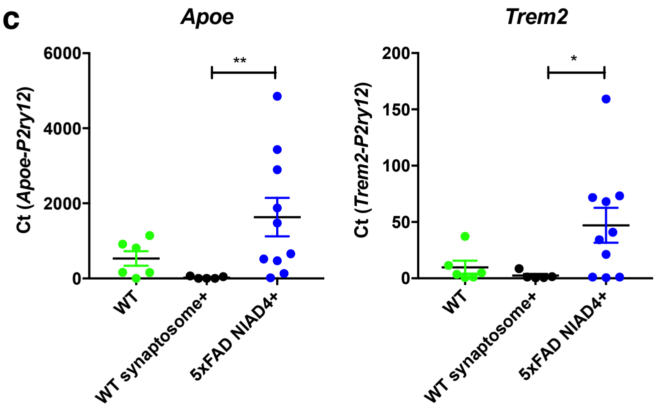

**Extended data Fig. 7**, Heatmap of gene expression in microglia isolated from OHSCs visualised with a k-nearest-neighbour graph rendered using a force directed layout[103], coloured by log2 transformed ΔCt values of selected genes upregulated in **a,** homeostatic microglia or **b,** NIAD4$^{+}$ (XO4$^{+}$) microglia. **c**, WT (*CX3CR1$^{GFP}$*) OHSCs were treated with pHrodo-Red-labelled synaptosomes, and synaptosome$^{+}$GFP$^{+}$ or synaptosome$^{-}$GFP$^{+}$ microglia were FACS-sorted 5 days later and profiled by qPCR for expression of the XO4$^{+}$ signature genes, *Apoe* and *Trem2*.

### Extended data Fig 8

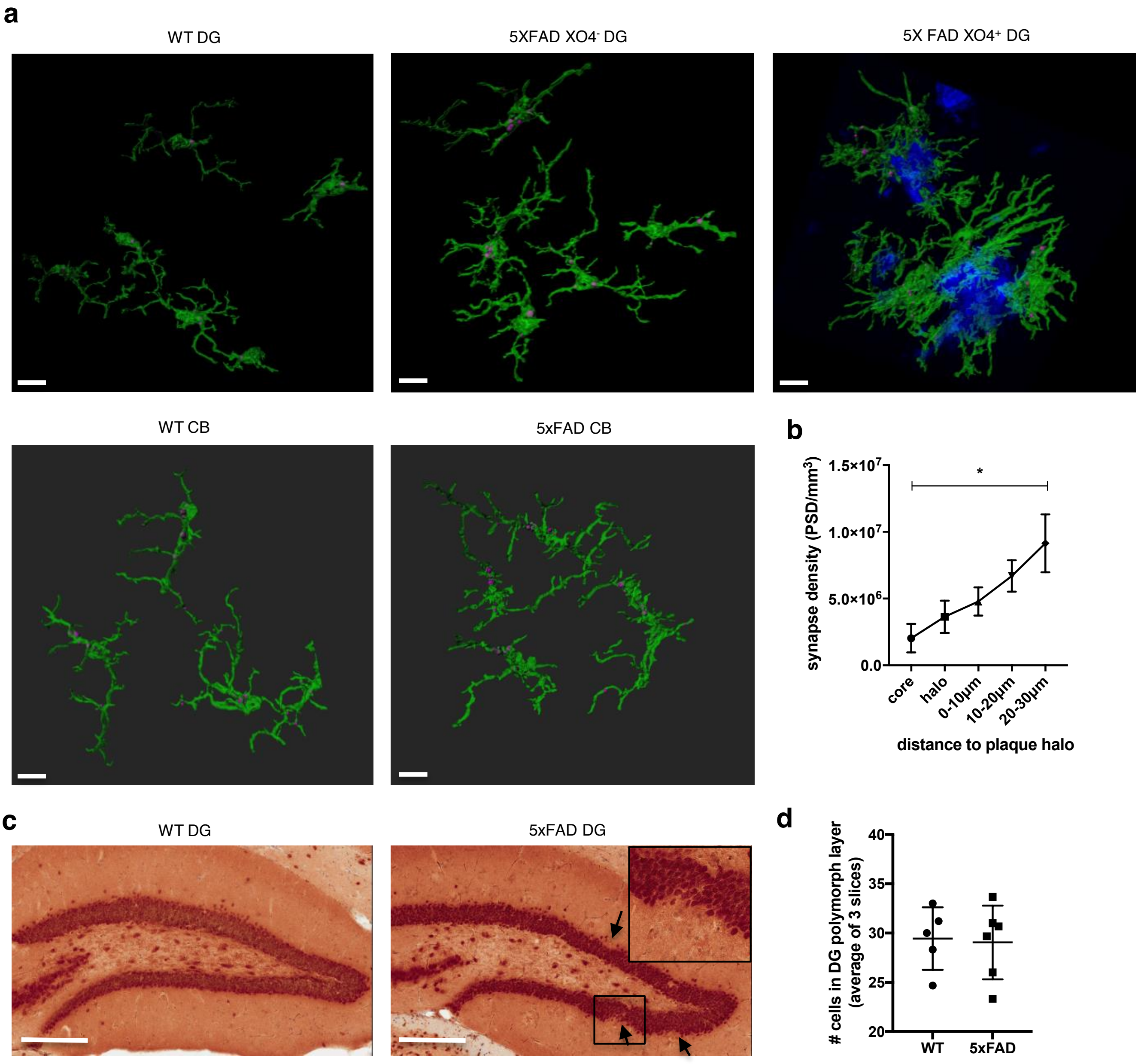


**e**

| | XO4+ vs 6M WT | | | | XO4+ vs XO4- | | | |
|---|---|---|---|---|---|---|---|---|
| **Gene** | **logFC** | **logCPM** | **PValue** | **fdr** | **logFC** | **logCPM** | **PValue** | **fdr** |
| ***C1d*** | -0.568 | 9.218 | 7.84E-04 | **2.40E-03** | -0.415 | 9.218 | 8.08E-02 | 1.83E-01 |
| ***C1qa*** | -0.319 | 13.586 | 5.03E-13 | **6.91E-12** | -0.463 | 13.586 | 2.27E-14 | **1.09E-12** |
| ***C1qb*** | -0.283 | 13.285 | 1.74E-11 | **2.04E-10** | -0.414 | 13.285 | 8.97E-13 | **3.51E-11** |
| ***C1qc*** | -0.404 | 13.159 | 6.84E-20 | **1.63E-18** | -0.476 | 13.159 | 4.03E-15 | **2.21E-13** |
| ***C2cd3*** | -0.168 | 9.059 | 3.82E-01 | 4.66E-01 | -0.688 | 9.059 | 6.65E-03 | **2.83E-02** |
| ***C3ar1*** | -0.215 | 9.381 | 1.68E-01 | 2.36E-01 | -0.672 | 9.381 | 7.63E-04 | **4.86E-03** |
| ***C5ar1*** | -0.923 | 9.284 | 3.70E-09 | **3.25E-08** | -0.791 | 9.284 | 2.75E-04 | **2.11E-03** |
| ***C5ar2*** | -0.916 | 9.063 | 3.55E-07 | **2.34E-06** | -0.734 | 9.063 | 2.17E-03 | **1.15E-02** |

**Extended data Fig. 8a**, Representative 3D reconstruction of confocal z-stacks showing PSD95 internalized within 6m WT, 5xFAD XO4$^-$ or 5xFAD XO4$^+$ microglia cells (scale bars = 15 μm). **b**, quantitation of synapse loss around plaques, measured by PSD95 puncta density (volume of PSD95 calculated using IMARIS spots function) in volumes of known distance in 10 μm increments from the plaque halo (0-30 μm). **c**, Immunohistochemistry showing NeuN labeling in the Dentate gyrus of 6m WT and 5xFAD mice and magnified inset. Arrows indicate altered localization of neuronal nuclei. **d,** neuron counts of the polymorph layer of dentate gyrus indicate no neuronal loss in this region, *n*=6 mice per genotype, and counts are averaged from 3 sagittal slices per mouse containing the hippocampal region. **e,** list of DE complement component and receptor genes in XO4$^+$ microglia compared to WT or XO4$^-$ microglia. FDR<0.05 is shown in bold.

### Extended data Fig 9

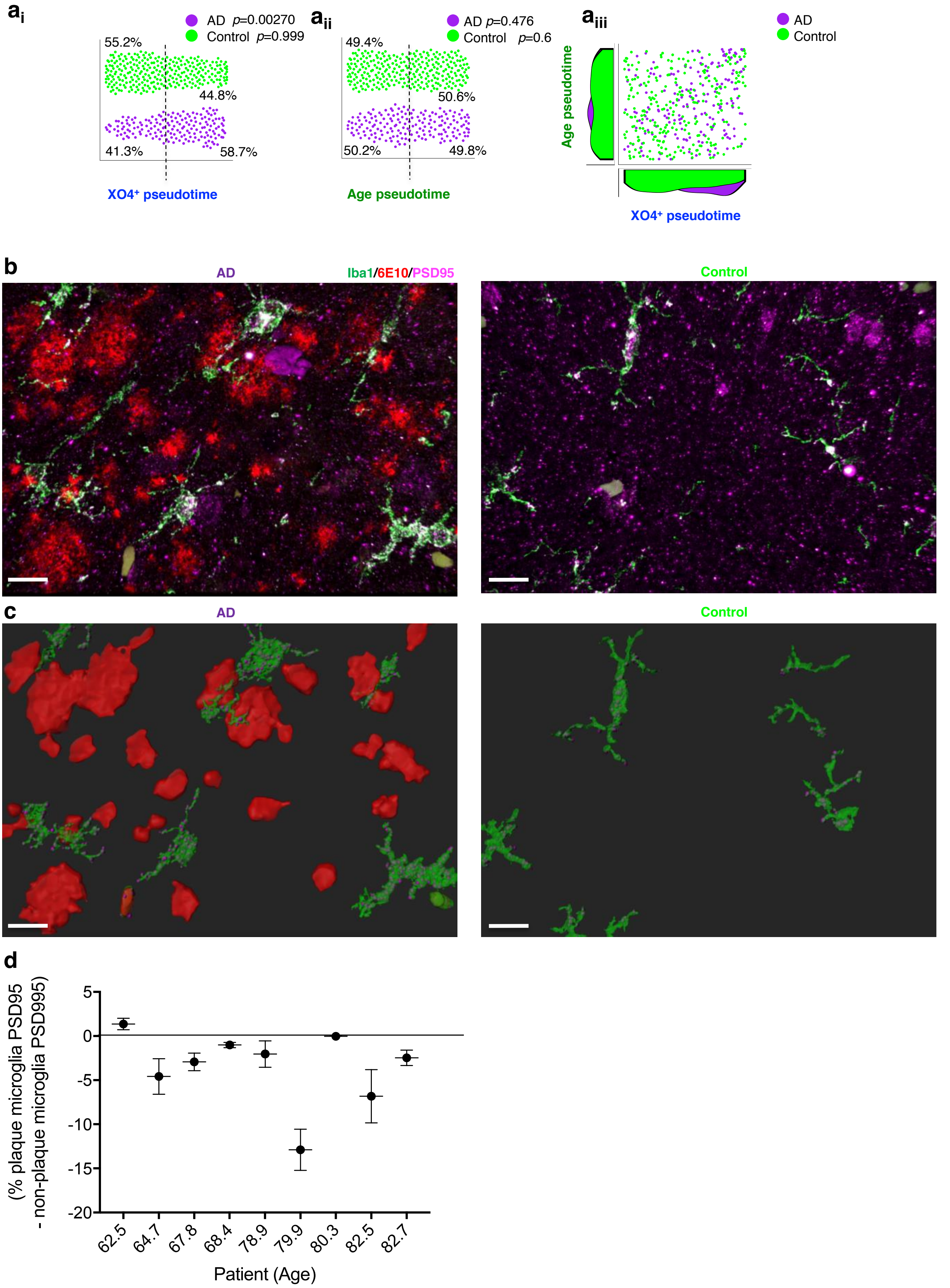

**Extended data Fig. 9a,** Diffusion maps pseudotime analysis of human control and AD microglial populations ordered by their expression of $XO4^+$ specific DEGs (**$a_i$**, taking top 10% of respective DE genes minus overlap with aging DEGs, 128 DEGs between 5xFAD $XO4^+$ and $XO4^-$ mice), ageing specific DEGs (**$a_{ii}$**, top 10% minus overlap with $XO4^+$ genes, 128 DEGs between 24M and 6M WT mice) and **$a_{iii}$**, scatter plot showing the relationship between ageing and $XO4^+$ pseudotime in individual cells and the density of cells at each point during the ageing (left) and $XO4^+$ (bottom) trajectories. **b,** Representative z-stack 3D projection and **c,** Imaris 3D reconstruction showing PSD95 internalized within microglia cell in human frontal cortex sections from AD patients ($n$=9) and cognitively normal individuals ($n$=8) stained with PSD95, 6E10 and Iba1 (scale bars = 15 µm)**. d,** quantitation of PSD95 internalized within microglia that are plaque adjacent or plaque distal. Data are mean ± SEM for individual microglia in each patient. $p$=0.02 using paired one-tailed one-sample $t$-test to test whether the mean differences in PSD95 within plaque adjacent compared to plaque-distal microglia are significantly different from 0.

### Extended data Fig 10

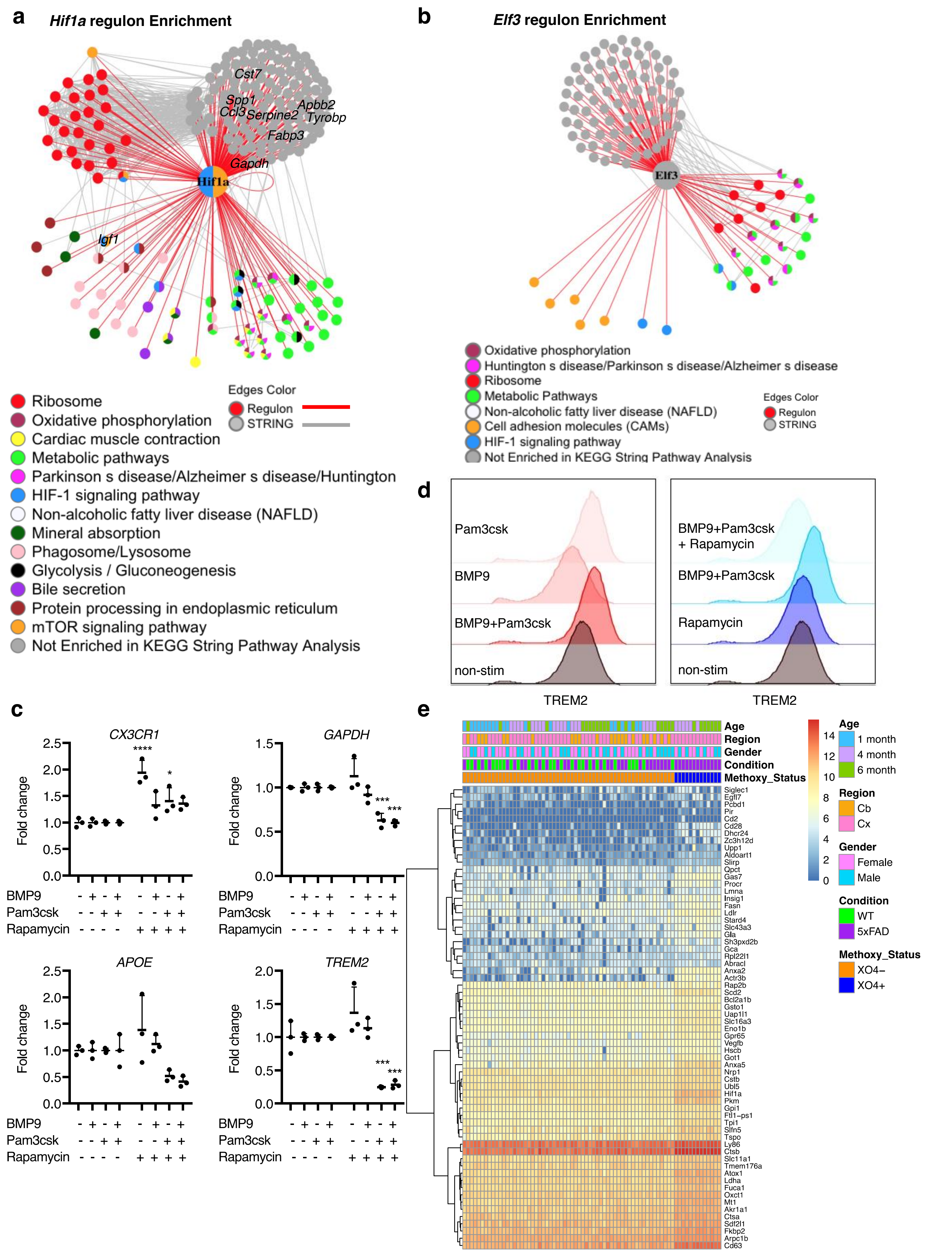

**Extended data Fig. 10a-b,** Network showing gene ontology enrichment of the genes in the *Hif1a* (**a**) and *Elf3* (**b**) regulons. Nodes are colored by their gene ontology(ies) and edges identified in STRING are grey, whereas those identified in our analysis are red. **c,** Stimulation of iMGLs with MyD88-dependent TLR agonist Pam3csk (alone or with BMP9) induces *GAPDH* as predicted, but also represses *APOE* and *TREM2*, $XO4^+$ signature genes not in the *Hif1a* regulon. Data are fold changes normalised to non-treated cells. **d**, Increased TREM2 surface protein expression in iMGLs following treatment with BMP9 and Pam3csk is reversed by the mTOR inhibitor, rapamycin, as measured by FACS. **e,** heatmap showing concordant overlap of DEGs in $XO4^+$ microglia from bulk RNA-seq data compared to human genes induced by Pam3csk and repressed by rapamycin in iMGLs.
